# Natural Variation in *Brachypodium distachyon* Responses to Combined Abiotic Stresses

**DOI:** 10.1101/2022.10.14.512283

**Authors:** Ella Ludwig, Seth Polydore, Jeffrey Berry, Joshua Sumner, Tracy Ficor, Erica Agnew, Kristina Haines, Kathleen Greenham, Noah Fahlgren, Todd C. Mockler, Malia A. Gehan

## Abstract

The growing world population increases demand for agricultural production, which is more challenging as climate change increases global temperature and causes more extreme weather events. High-throughput phenotyping tools can be used to measure plant responses to the environment to identify genomic regions associated with response to stress. This study examines the phenotypic variation of 149 accessions of *Brachypodium distachyon* under drought, heat, and the combination of both stresses. Heat alone causes the largest amounts of tissue damage and the combination of heat and drought causes the largest decrease in plant biomass compared to other treatments. Notably, Bd21-0, the reference line for *B. distachyon*, was identified as not having very robust growth under stress conditions, especially in the heat-drought combined treatment. Climate data from the collection locations of these accessions (climate of origin) was used to assess whether climate of origin was correlated with responses to stresses and it was found to be significantly associated with height and percent of plant tissue damage. Additionally, genome wide association mapping found a number of genetic loci associated with changes in plant height, biomass, and the amount of damaged tissue under stress. Some SNPs found to be significantly associated with a response to heat or drought are also significantly associated in the combination of stresses, while others are not, and some significantly associated SNPs were only identified in the combined stress treatment. This, combined with the phenotypic data, indicates that the effects of these abiotic stresses are not simply additive, and the responses of *B. distachyon* to the combined stresses differ from drought and heat alone. Significant SNPs were closely located to genes known to be involved in plant responses to abiotic stresses.

## Introduction

The increasing global population is likely to create a number of challenges in the coming decades. The world population is expected to reach 9.8 billion people by the year 2050, which means there will be nearly 2 billion additional people on the planet to feed in fewer than 30 years (Nations, 2019). With more than a billion people in the world currently experiencing some form of food insecurity or undernourishment, the increasing global population creates an even higher demand for food, fuel, and fiber, that current production levels cannot meet (Conway, 2012; Swaminathan, 2012; FAO et al., 2020; Roy et al., 2006).

An additional complication is that future crops will need to withstand the changing climate. Due to human activity, greenhouse gas concentrations in the atmosphere are rising, which is causing significant changes in global temperatures and overall climate patterns (IPCC, 2019). Global temperatures are expected to rise by 3-5°C in the next 50-100 years, and experts expect to see an increased frequency of extreme weather events like drought and high temperatures (IPCC, 2014, 2019). Therefore, crops need to be more productive under more challenging environments to keep up with global demand.

Environmental stress tolerance is a key factor for crop productivity, especially in a changing climate, as abiotic stress is a major cause of reduced crop yields across the globe (Atkinson and Urwin, 2012). Abiotic stresses are estimated to reduce average yields by more than 50% for many major crops since they can have an adverse impact on nearly all stages of plant growth and development, including leaf damage, accelerated leaf senescence, reduced photosynthetic capacity, and can greatly reduce plant biomass and grain production and therefore economic yield (Allakhverdiev et al., 2008; Farooq et al., 2011; Atkinson and Urwin, 2012). This means that there will be an ever-growing demand for stress-tolerant varieties as climate change brings with it more of these biomass- and yield-reducing stresses. To reverse this trend, it is important to study plant responses to abiotic stresses.

Historically, single abiotic stresses are more often studied than stresses in combination (Rasmussen et al., 2013; Atkinson and Urwin, 2012; Rizhsky et al., 2004). However, plant response to combinations of abiotic stress can result in transcriptional changes that differ from the simple summation of each individual stress (Rasmussen et al., 2013; Rizhsky et al., 2002, 2004). In some regards it is not surprising that plants have distinct responses to combined stress conditions in comparison to individual stresses since many frequently co-occurring stresses can result in contradicting plant responses (Mittler, 2006; Anderson et al., 2004; Asselbergh et al., 2008; Atkinson and Urwin, 2012; Mittler and Blumwald, 2010; Balfagón et al., 2020; Suzuki et al., 2016; Choudhury et al., 2017). For example, a common response by plants under drought stress is closing their stomata to preserve water, but in heat stress they would open them to cool their leaves to prevent tissue damage (Rizhsky et al., 2004). This means that in the combination of both drought and heat stresses, plant responses could look vastly different than when exposed to just one stress alone. Therefore, it is imperative to study not only plant responses to individual stresses but also responses to combinations of stresses to work towards making new crop varieties. Since many major crops have been bred for high yield but not necessarily for abiotic stress tolerance, it is valuable to explore and exploit the natural diversity of stress tolerance in weedy relatives to current elite crops (Mickelbart et al., 2015).

*Brachypodium distachyon* is a weedy Pooid C_3_ grass that is closely related to important monocot food crop species including wheat, barley, and rice (International Brachypodium Initiative, 2010). *B. distachyon* also has very similar cell-wall composition and overall architecture to major bioenergy grasses such as miscanthus and switchgrass (Gomez et al., 2008; Coomey and Hazen, 2015). In addition to these genetic or morphological similarities to major food and bioenergy crops, *B. distachyon* has a small stature, short generation time, is easy to cultivate and transform, and has genomic resources available, which all combine to make it a very powerful and attractive model plant (Mur et al., 2011; Brutnell et al., 2015; Kellogg, 2015). *B. distachyon* accessions have been collected throughout its native range in the Mediterranean and Middle East (Draper et al., 2001; Garvin et al., 2008; Filiz et al., 2009; Tyler et al., 2016). As is the case for many species, the reference accession of *B. distachyon* (Bd21-0), is not necessarily one with ideal responses to abiotic stress conditions, and relatively few studies have examined the phenotypic variation in other *B. distachyon* accessions under the same conditions (Luo et al., 2011; Colton-Gagnon et al., 2014; Shi et al., 2015; Rivera-Contreras et al., 2016; Jiang et al., 2017; Des Marais et al., 2017; Benavente et al., 2013; Chen and Li, 2016; Cao et al., 2016b, 2016a; Liu and Chu, 2015; You et al., 2015). Even fewer studies have examined *B. distachyon* accessions under combinations of stress, which, as mentioned earlier, is necessary to get a more representative picture of how *B. distachyon* would respond to stress in the field or in the wild (Des Marais et al., 2017).

A review of *B. distachyon* literature demonstrates additional challenges in studying abiotic stress, and more specifically drought, as the same accessions are conflictingly described as drought tolerant or drought susceptible in different studies. In 2011, Luo et al. classified the *B. distachyon* reference accession, Bd21-0, as susceptible to drought stress compared to 56 others, taking into account leaf water content, wilting, and chlorophyll fluorescence (Luo et al., 2011). The same study also described Bd1-1 as a relatively drought tolerant accession, using the same measured traits (Luo et al., 2011). Shi et al., on the other hand, identified Bd21-0 as being tolerant to drought when examining survival, chlorophyll content, and electrolyte leakage (measurement of membrane damage) as measures of drought tolerance (Shi et al., 2015). Des Marais et al. examined both heat and drought stress and their combinations and identified Bd1-1 to be one of the lowest yielding accessions under all combinations of drought and heat conditions tested (Des Marais et al., 2017). Less is known about the heat tolerance of these lines, but Des Marais et al. found that Bd21-0 had higher seed yield in hot but ample water conditions (Des Marais et al., 2017).

Conflicting results on the same accessions underline the difficulty in studying drought, and could be partially due to the variety of ways in which drought can be applied and defined in experiments. Experimental application of water-limitation (drought) can differ greatly, which complicates the ability to directly compare results between studies. Acute drought stress can be applied in a laboratory setting by completely removing a plant from soil or growth media (Legnaioli et al., 2009; Gagné-Bourque et al., 2015). Progressive drought treatments, on the other hand, can be applied by maintaining reduced watering levels compared to a control level over a longer period of time (Fahlgren et al., 2015; Granier et al., 2006; Skirycz et al., 2011). In a different kind of progressive drought treatment, water is withheld, allowing the soil to dry over time (Des Marais et al., 2017; Jiang et al., 2017; Luo et al., 2011; Shi et al., 2015). Progressive drought treatments are generally more representative of drought in the field than acute drought stress treatments are, but can often be more challenging to reproduce, since manually maintaining a specific reduced level of soil moisture is labor-intensive, and the severity of the drought stress depends on the consistency of soil drying between replicates and experiments (Zhang et al., 2004; Sun et al., 2014; Claeys and Inzé, 2013).

Similarly, there are diverse ways to apply heat stress to plants, which makes it difficult to compare results across different experiments. Acute heat stress is often applied by exposing plants to high temperatures for a relatively short period of time (generally a few hours) and then taking measurements for traits of interest directly after the heat treatment (Sun et al., 2017). Progressive heat treatments are often performed by maintaining high ambient temperatures for the entire course of the experiment after germination (Fahlgren et al., 2015; Des Marais et al., 2017; Hedhly et al., 2020). Alternatively, exposing plants to high heat every day for a certain length of time around midday can be used to simulate a common form of progressive heat stress in the field (Samakovli et al., 2020).

To address these issues of inconsistency, reproducibility, and high labor input requirements in implementing abiotic stresses, automated watering and growth systems have been developed. Within the last decade, these automated systems in greenhouses, growth chambers, and experimental fields have advanced considerably making it possible to apply abiotic stress treatments reproducibly at population scale (Chen et al., 2014; Fahlgren et al., 2015; Granier et al., 2006; Skirycz et al., 2011). This study uses an automated watering and image-capture system to apply combinations of drought and heat stress to a diverse population of *B. distachyon* accessions in two experiments. Since imaging is a non-destructive measurement, assessing stress through image data allows the progression of stress to be tracked over time. To analyze image data from these two phenomics experiments, this project utilizes high-throughput image analysis tools. The field of image-based high-throughput phenomics develops methods to capture, extract, and utilize plant trait information over time from image data. PlantCV is an open-source, open-development image analysis package written in Python that enables users to develop image processing workflows, then parallelize those workflows over large sets of image data (Gehan et al., 2017; Fahlgren et al., 2015). This software has been used to extract numerical data from images to estimate various traits including biomass (Fahlgren et al., 2015) and plant health (Enders et al., 2019; Zheng et al., 2019) for various plant species (Gehan et al., 2017).

Numerical trait information extracted from image data, along with low coverage sequencing data, is used to carry out a genome wide association study (GWAS) on 148 accessions of *B. distachyon* accessions. A GWAS is a way to inspect the genome for genetic polymorphisms, usually single nucleotide polymorphisms (SNPs) that are found more frequently in organisms with the trait being assessed (Ingvarsson and Street, 2011; Wilson et al., 2015). GWA studies can be used to explore the genetic architecture of complex traits in plants, identifying the number of alleles, genetic loci, and effect sizes, associated with a certain phenotype (Ingvarsson and Street, 2011; Wilson et al., 2015). Complex traits would normally be difficult to examine without GWAS, since most adaptive traits are influenced by many genes that each individually have a small to moderate effect on the phenotype (Ingvarsson and Street, 2011). This is true for drought and heat stresses, as plant responses to these abiotic stresses have been shown to be polygenic and not as easily isolated and identified as other traits like flowering time (Tyler et al., 2016). Thousands of GWAS have been conducted in major crops such as rice, wheat, maize, and sorghum, which have included a variety of traits examining everything from yield to plant development to characteristics associated with biotic and abiotic stresses (Gupta et al., 2019). Though currently often underutilized, the results of GWAS can be used for crop improvement (Gupta et al., 2014). Genetic loci of interest identified in a GWAS can be further examined and then used to develop improved cultivars through traditional breeding methods or through genetic engineering (Gupta et al., 2019). One important documented example of the use of GWAS results to improve a major crop was the development of a line of maize with increased pro-vitamin A content (Xiao et al., 2017).

There have been a few GWAS done in *B. distachyon*, but compared to major crop species and other important model plants, the genetic architecture of *B. distachyon* remains relatively unexplored (Dell’Acqua et al., 2014; Tyler et al., 2016; Wilson et al., 2015; Lee, 2016; Wilson et al., 2019). Multiple studies have identified genetic loci associated with flowering time in. *B. distachyon* (Wilson et al., 2019; Tyler et al., 2016), others have explored genetic loci associated with environmental adaptation or fitness in their different climates of origin (Wilson et al., 2015; Dell’Acqua et al., 2014), and genetic loci associated with biofuel-related traits such as biomass accumulation have also been identified (Wilson et al., 2019; Lee, 2016).

In this study, the population diversity and natural variation of this *B. distachyon* population under combinations of progressive heat and drought stresses are examined. These accessions were collected throughout the native growth range of *B. distachyon*, and come from diverse climates of origin with a wide range of elevations, temperatures, and precipitation patterns (Figure 1, Supplemental Figure 1). At this time, this is the largest temporal phenotypic assessment of this crucial model plant to date and also provides the community with additional genotype information because 132 accessions do not overlap with the accessions genotyped in Tyler et al. (Tyler et al., 2016). A GWAS was performed to investigate the underlying genetic architecture in *B. distachyon* that is associated with changes in plant phenotype in each of the drought and heat stress conditions, which will also provide new information about the *B. distachyon* genome and its involvement in abiotic stress responses. This study examines *B. distachyon* as a model for bioenergy grasses, so the focus traits are biomass accumulation, height, and percent of healthy photosynthetically active tissue.

**Figure 1.**
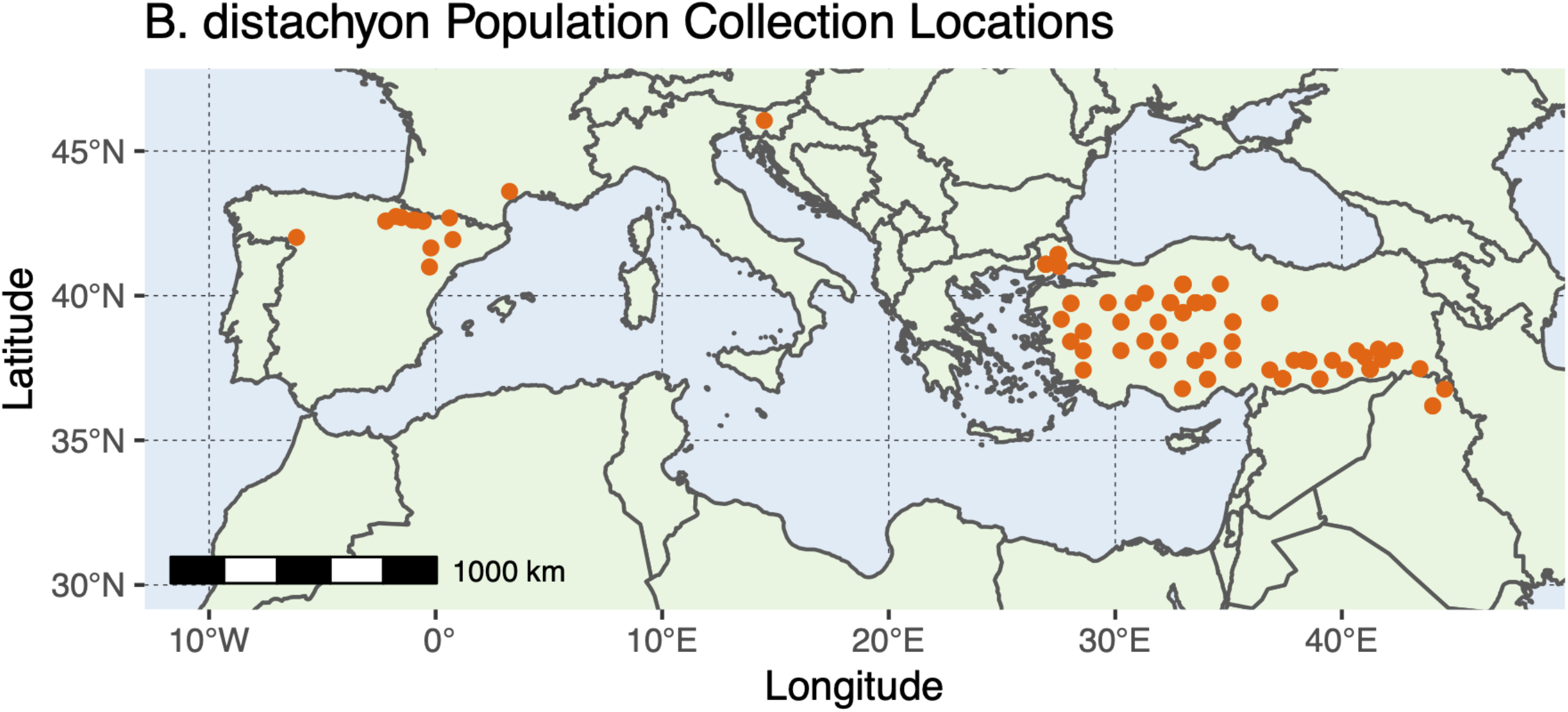
(collection location map): Map of Southern Europe, Northern Africa and the Western Middle East showing the distribution of collection locations of 147 of the *B. distachyon* accessions included in the analysis of the data collected in this experiment (specific collection location is unavailable for two of the 149 accessions). The x- and y-axes represent longitude and latitude on the Earth’s surface, respectively. Scale bar in the bottom left represents 1000 km.

## Results

### Experimental Overview

To examine the natural variation of *B. distachyon* under drought, heat, and combined drought and heat, low-coverage sequencing data (genotyping-by-sequencing; GBS) and two high-throughput phenotyping experiments using the Bellwether Phenotyping Platform at the Donald Danforth Plant Science Center (Fahlgren et al., 2015) were done. In the first phenotyping experiment (Figure 2A), 137 *B. distachyon* accessions were grown under control or water-limited conditions (20% of control watering, starting on day 5 on the phenotyping platform). In the second experiment (Figure 2B), an overlapping population of 144 *B. distachyon* accessions were treated with heat or heat and water-limited conditions (20% of control watering, starting on day 5 on the phenotyping platform). Because the Bellwether Platform growth chamber is not subdivided to accommodate two different temperature conditions, the two experimental treatments could not be done simultaneously. Therefore, the resulting images from these two experiments were processed independently but the normalized outputs were compared. Later portions of the Materials and Methods section provide a more detailed description of the analysis methods. Over the course of both experiments, more than 185,000 RGB images were collected. To extract numerical information from these images to estimate plant health and shape parameters, this project utilizes high-throughput plant phenomics software PlantCV (Fahlgren et al., 2015; Gehan et al., 2017), the details of these analyses are in the Materials and Methods section.

**Figure 2.**
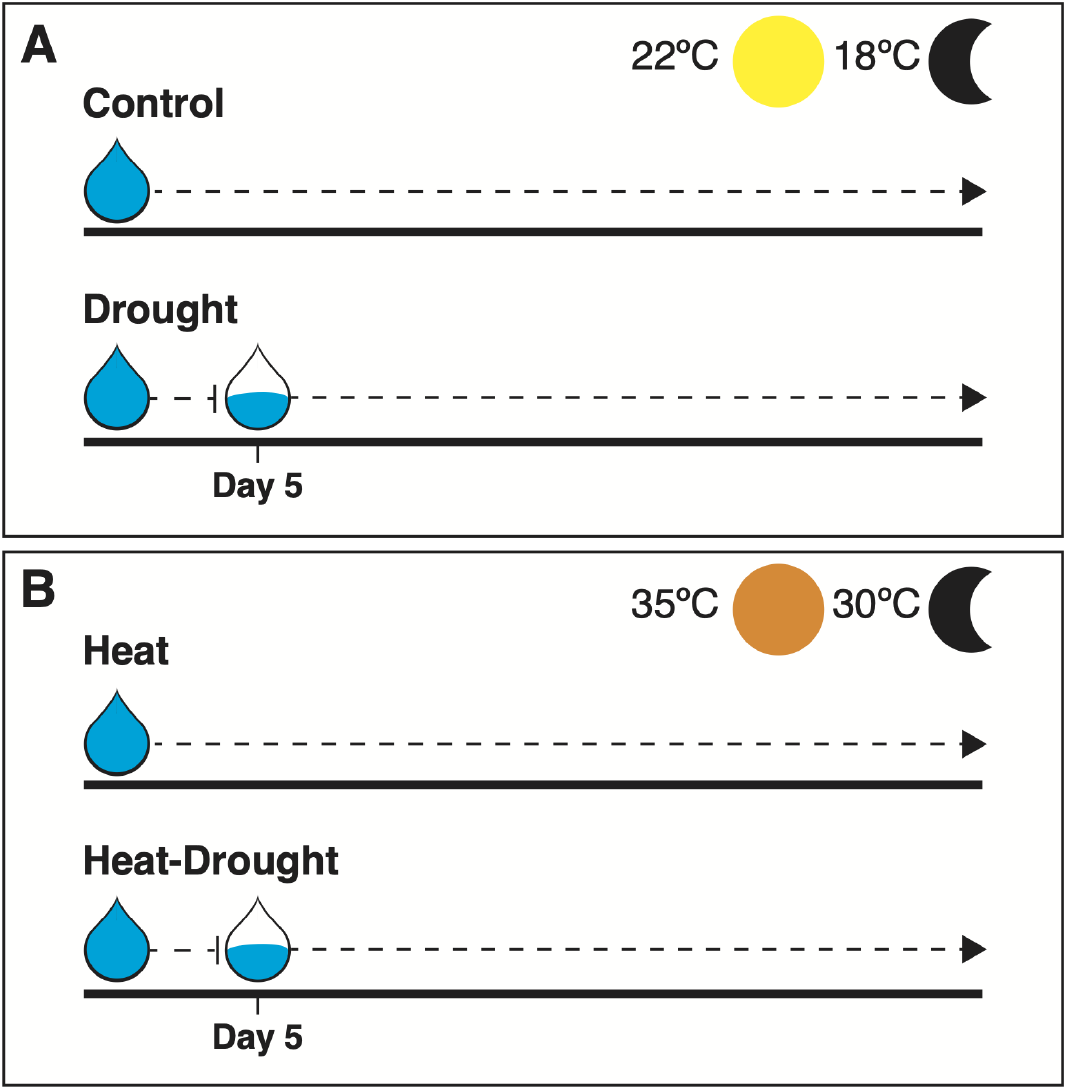
(experimental design): Experimental conditions for both experiments conducted. A) Experiment with control and drought treatments. Both were grown at 22°C during the days and 18°C during the nights, with the drought treated plants receiving 20% of control watering starting on day 5 on the phenotyping platform. B) Experiment with heat and combination of heat and drought treatments. Both were grown at 35°C during the day and 30°C during the night, with heat and drought combination treated plants receiving 20% of control watering starting on day 5 on the phenotyping platform. All plants were grown under a 14h photoperiod.

### Abiotic stress treatments can be differentiated by PCA over time

A principal component analysis (PCA) was used on trait data extracted from images with PlantCV (percent_unhealthy, area, height_above_reference, hue_circular_mean, width, height, convex_hull_area, solidity, perimeter, longest_path, ellipse_major_axis, ellipse_minor_axis, ellipse_angle, ellipse_eccentricity) to assess the effects of the different treatment conditions in this study (Figure 3). The PCA analysis shows overlap of all experimental conditions on Imaging Day 1 prior to the application of different treatments, suggesting that despite the separate experiments, no substantial phenotypic differences between plants were discernible. Over the subsequent days of the experiment, the different treatment groups begin to separate based on their phenotypic differences, and by the final day of the experiment the PCA results show strong separation between most treatments (Figure 3). It is interesting to note that there is a strong separation between the drought and heat treatments as well as drought and heat-drought stresses, but there is substantial overlap between the heat and heat-drought treatment groups (Figure 3).

**Figure 3.**
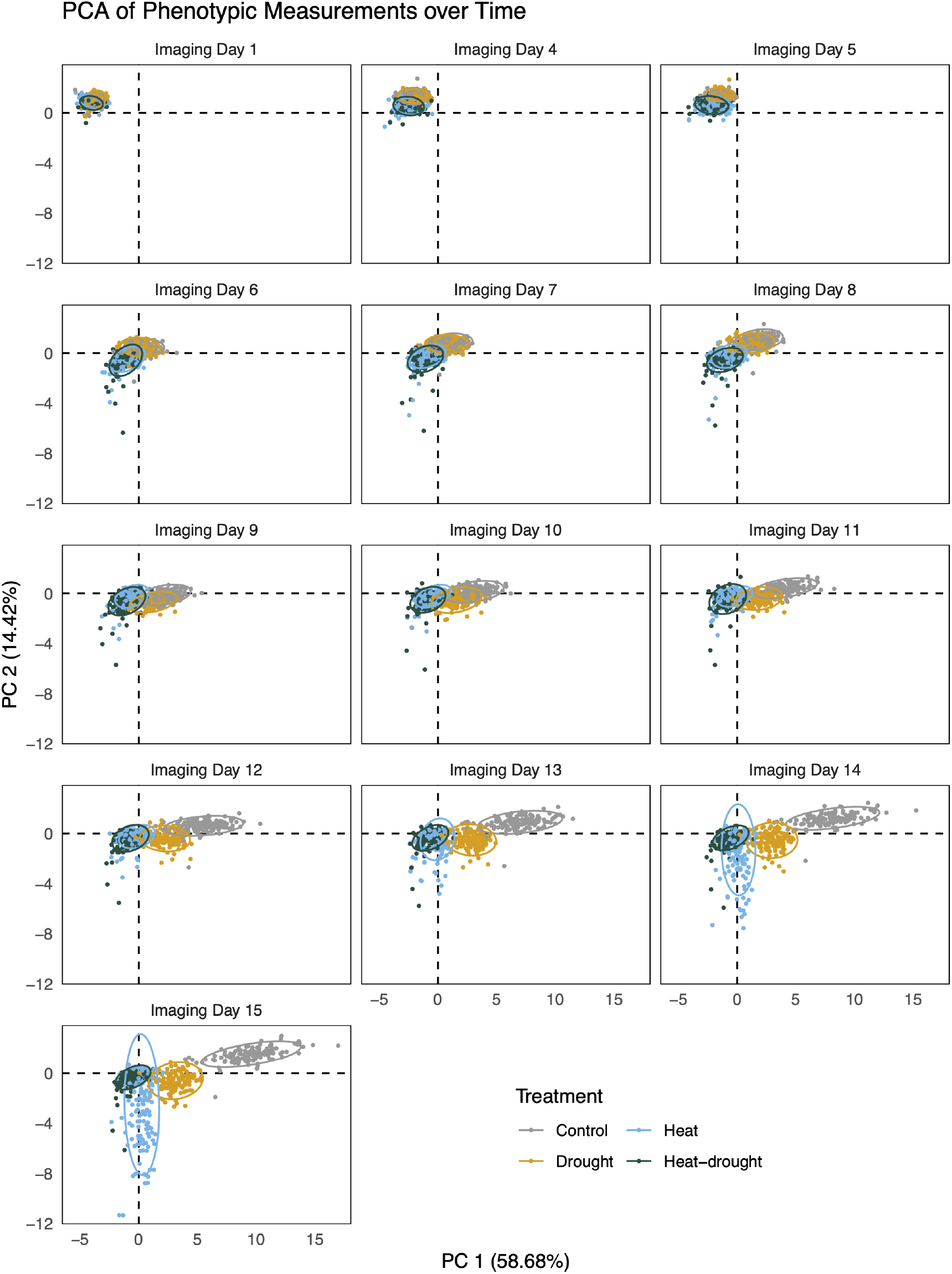
(phenotype PCA): Principal component analysis of *B. distachyon* responses to different experimental conditions over time using trait data extracted from images with PlantCV Traits included are: percent_unhealthy, area, height_above_reference, hue_circular_mean, width, height, convex_hull_area, solidity, perimeter, longest_path, ellipse_major_axis, ellipse_minor_axis, ellipse_angle, ellipse_eccentricity. The graph shows principal component 1 (PC 1) on the x-axis, which explains 58.68% of the variance, and principal component 2 (PC 2) on the y-axis, which explains 14.42% of the variance, with 95% confidence ellipses included. Points are colored by treatment.

### Heat stress alone resulted in the most tissue damage in comparison to drought and the combined stress

The naive Bayes classifier from PlantCV calculates the statistical likelihood that a pixel belongs into user defined classes (Abbasi and Fahlgren, 2016; Gehan et al., 2017). To estimate the effects of individual and combined drought and heat stresses on tissue damage in *B. distachyon*, two classes for healthy and unhealthy tissue were defined. The amount of unhealthy versus healthy plant tissue was then calculated as a percentage of total plant shoot area identified and categorized as “percent damage”. When examining these data across the *B. distachyon* population, it is clear that there is considerable variation in response to the different stress treatments (Figure 4). Some accessions, such as BdTR3M, have relatively low percentages of damage across all four conditions, while others, such as BdTR2B and BdTR2G, have relatively high percentages of damage across conditions. A number of accessions also have high percentages of unhealthy tissue in some stress treatments but not in others. For example, Adi-4 and BdTR5E have among the highest percent damage in heat and heat-drought, but among the lowest in the drought treatment, whereas BdTR3R and Tek-5 have among the highest percent damage in the drought and heat-drought treatments, but among the lowest in the heat treatment. An unexpected result when looking at the percent damage on these plants was that heat stress alone seems to have the most estimated leaf damage in comparison to drought, or the combination of stresses (Figure 4). The Naive Bayes classifier data showing this is corroborated when visually examining the images of these plants (Figure 5). Bd21-0, often used as a reference line for *B. distachyon*, has more than double the percent damage in heat compared to both drought and the heat-drought combination (Supplemental Figure 2).

**Figure 4.**
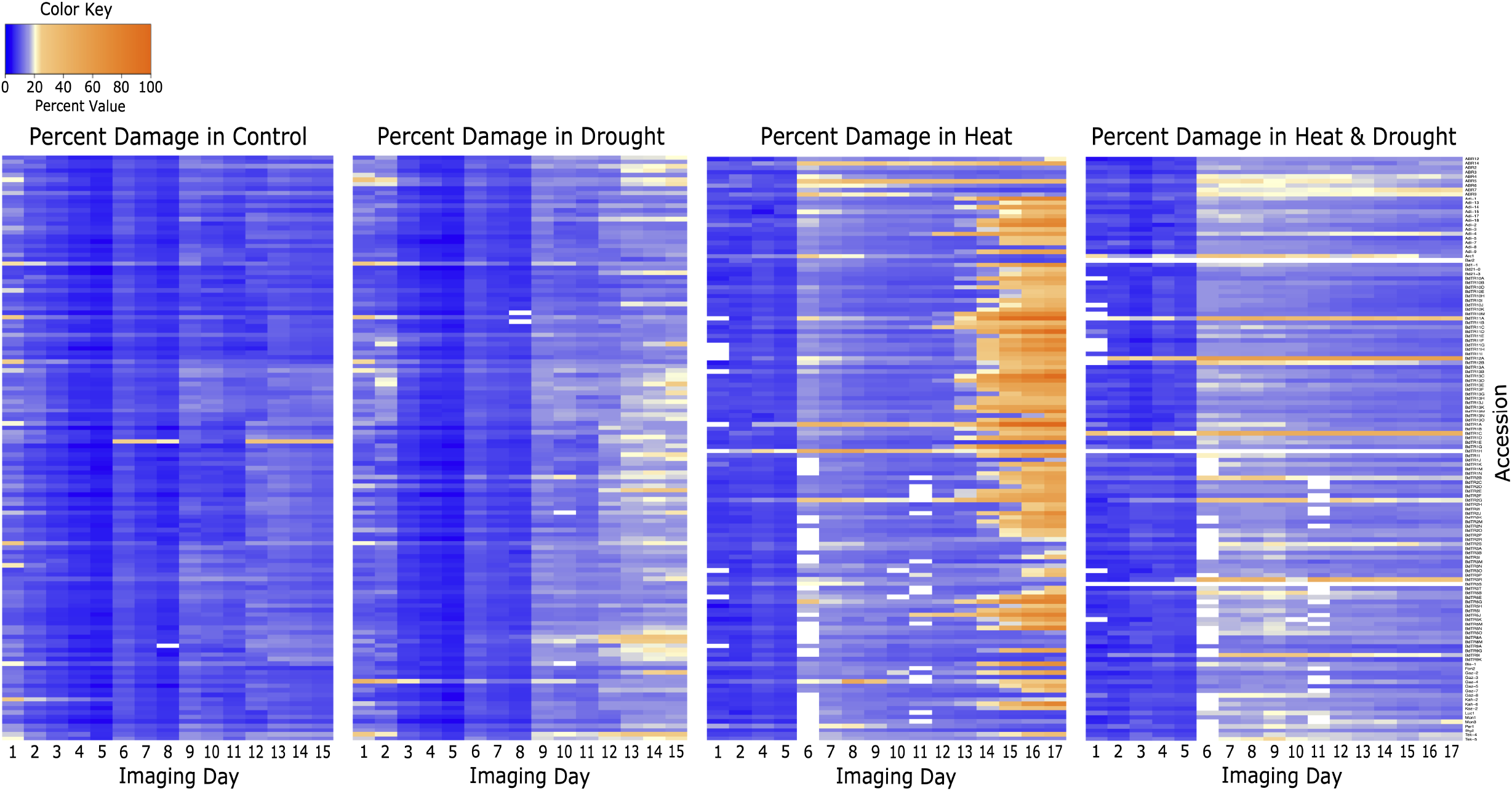
(percent damage heatmaps): Heatmaps of the percentage of unhealthy tissue in plants from the naive Bayes classifier, in the four experimental conditions (from left to right: control, drought, heat, heat & drought combined). The x-axis of each heatmap shows “Imaging Day,” which represents each imaging cycle, since half of the plants were imaged on even days and half on odd (so Imaging Day 1 is all images taken on day 1 *and* day 2 of the experiment). Imaging Day 3 was removed from the heat treatment. On the y-axis are the accessions, sorted alphabetically in descending order.

**Figure 5.** (panel of lines): Panels showing images of plants from two accessions in all four treatments. The top, outlined in orange, shows accession BdTR11F, which has a very high percentage of its tissue classified as unhealthy in heat, and the bottom, outlined in blue, shows accession Adi-4, which has a very low percentage of its tissue classified as unhealthy in heat.

### The combination of heat and drought stresses resulted in the greatest decrease in biomass compared to heat or drought alone

PlantCV outputs were also used to examine biomass accumulation of *B. distachyon* in these abiotic stress treatments by assessing both plant height and plant shoot area. Heatmaps of plant height show that while there is variation across the population, the majority of accessions are still able to grow relatively tall in the drought treatment compared to the other stresses, especially in comparison to the combination of heat and drought (Figure 6). A similar pattern is seen for plant shoot area across treatments and accessions (Supplemental Figure 3). Bd21-0 follows this trend, and is among the top accessions for area and height in the drought treatment, but is one of the smallest accessions in the heat and heat-drought treatments (Supplemental Figure 2). Two accessions, BdTR11B and Mon3, are among the top accessions for both area and height across all four experimental conditions (Supplemental Figure 2), showing robust growth across stress treatments. Conversely, accessions such as BdTR5G and ABR12 were among the bottom of all accessions across stress treatments for area and height (Supplemental Figure 2). Overall, the combination of drought and heat stresses seem to have a more detrimental impact on biomass accumulation than heat or drought stress alone, though heat alone is a close second (Figure 6, Supplemental Figure 3).

**Figure 6.**
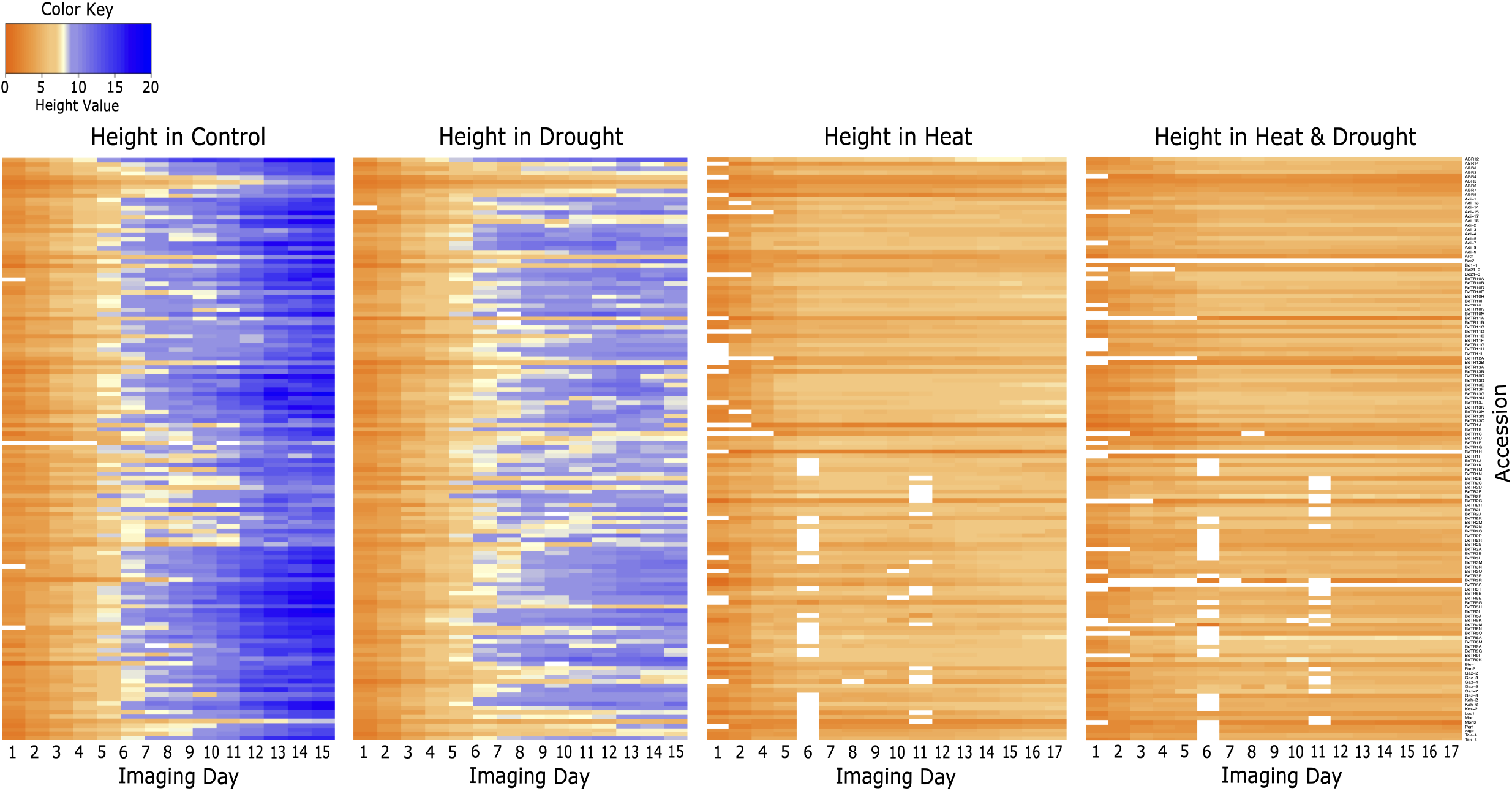
(height heatmaps): Heatmaps of the average plant height in the four experimental conditions (from left to right: control, drought, heat, heat & drought combined). This was calculated by taking the mean of the height of all plants of the same accession in the same treatment in each imaging cycle. The x-axis of each heatmap shows “Imaging Day,” which represents each imaging cycle, since half of the plants were imaged on even days and half on odd (so Imaging Day 1 is all images taken on day 1 *and* day 2 of the experiment). Imaging Day 2 was removed from the control and drought conditions, and Imaging Day 3 was removed from the heat treatment. On the y-axis are the accessions, sorted alphabetically in descending order.

This suggests that *B. distachyon* has non-overlapping responses that decrease biomass accumulation in response to the combination of drought and heat stress in comparison to individual stresses. Additionally, these data seem to indicate that heat is the more severe stress compared to drought, since the plants were generally able to maintain biomass in the drought treatment but were considerably smaller in both the heat and heat-drought treatments. However, the results from the naive Bayes classification would suggest that although plants are smaller under the combined stress conditions there is a lower percentage of damaged tissue, and therefore a higher amount of photosynthetically active tissue under the combined stress in comparison to heat stress alone.

### Identification of stress-tolerant and stress-susceptible accessions

To assess the natural diversity in *B. distachyon* resilience to drought and heat stresses, the difference was taken between each accession’s phenotype in each of the three stress treatments (drought, heat, heat and drought combination) and their control for the three main phenotypic traits being assessed in this study: height, area, and percent damage. Accessions with the smallest deficits in growth and health across the stresses compared to control conditions (not only in one stress) were identified as having resilient growth under stress, while accessions with the biggest decreases in size and healthy tissue under stress conditions compared to control were identified as not having resilient growth under these stresses. The two accessions identified as being most resilient to heat and drought stress conditions are BdTR1E, which was in the top 25% of all accessions for the three phenotypes assessed in all stress treatments, and BdTR10E, which was in the top 25% of all accessions for all phenotypes and stresses except for percent damage in the heat treatment, where it was in the top 50% (Supplemental Figure 2). The two accessions identified as being the overall least resilient to these stresses are Adi-18, which was in the bottom 25% of accessions for all phenotypes and treatments except for percent damage in heat, where it was in the bottom 50%, and BdTR5J, which was in the bottom 25% for all traits and treatments except the percent damage in the drought and heat-drought treatments, where it was in the bottom 50% and middle 50%, respectively (Supplemental Figure 2).

A similar methodology was used to identify accessions that are tolerant or susceptible to a single stress treatment. Notable results were that BdTR1E performs well in all three stress treatments (as previously mentioned), and seems to have mechanisms to maintain healthy, photosynthetically active tissue with smaller size reductions under these stress conditions compared to the other accessions included in this study (Supplemental Figure 4). This could be an interesting accession to examine in future studies. Another noteworthy result is that Bd21-0, a commonly used reference line for *B. distachyon*, performed very poorly in the heat-drought combination stress treatment, and was in the bottom 25% of all accessions in each of the three stress treatments (Supplemental Figure 4). Bd21-0 is below average for some phenotypic traits and near average for others in the drought and heat stresses alone, but is also never among the best performing accessions. Complete lists of the accessions identified as being tolerant or susceptible to individual stresses can be found in the Supplemental materials (Supplemental Figure 4).

### Treatment effect is stronger and appears earlier in heat and heat-drought treatments than in drought

For each stress treatment compared to control, the partial correlations, or percent variance explained, of genotype, treatment and the genotype-treatment interaction (GxE) were calculated using a variance components model. The analyses were done using data from four time points throughout the experiments, corresponding to Imaging Days 4, 7, 10, and 15 (the final day with data for all treatments), to assess the variance explained over time. The variance explained was plotted in stacked bar graphs to show both the total variance explained for each phenotype measured as well as the amount explained by genotype, treatment and the genotype-treatment interaction, in addition to the amount unexplained (Figure 7). Overall, the genotype and genotype-treatment interaction components were largest at Time Point 1 and decreased over time as the treatment effect increased (Figure 7). This is especially true for the drought treatment, where the treatment component only begins to account for any meaningful portion of the variance explained in Time Point 2. In the heat and heat-drought treatments, however, the treatment component accounts for a much larger portion of the variance explained earlier, with this component making up over 70% of the variance explained for some variables already at Time Point 1 in both the heat and heat-drought treatments, compared to less than 1% in the drought treatment. Additionally, the variance explained by the treatment is larger overall in the heat and heat-drought treatments compared to the drought treatment. The largest is 89.37% and 89.96% for any variable – in the heat and heat-drought treatments, respectively, at Time Point 4, compared to 79.26% in the drought treatment (Figure 7). Since both treatments that involve heat (heat and heat-drought) follow this trend, in contrast to the drought treatment, this indicates that the heat treatment is the more severe stress.

**Figure 7.**
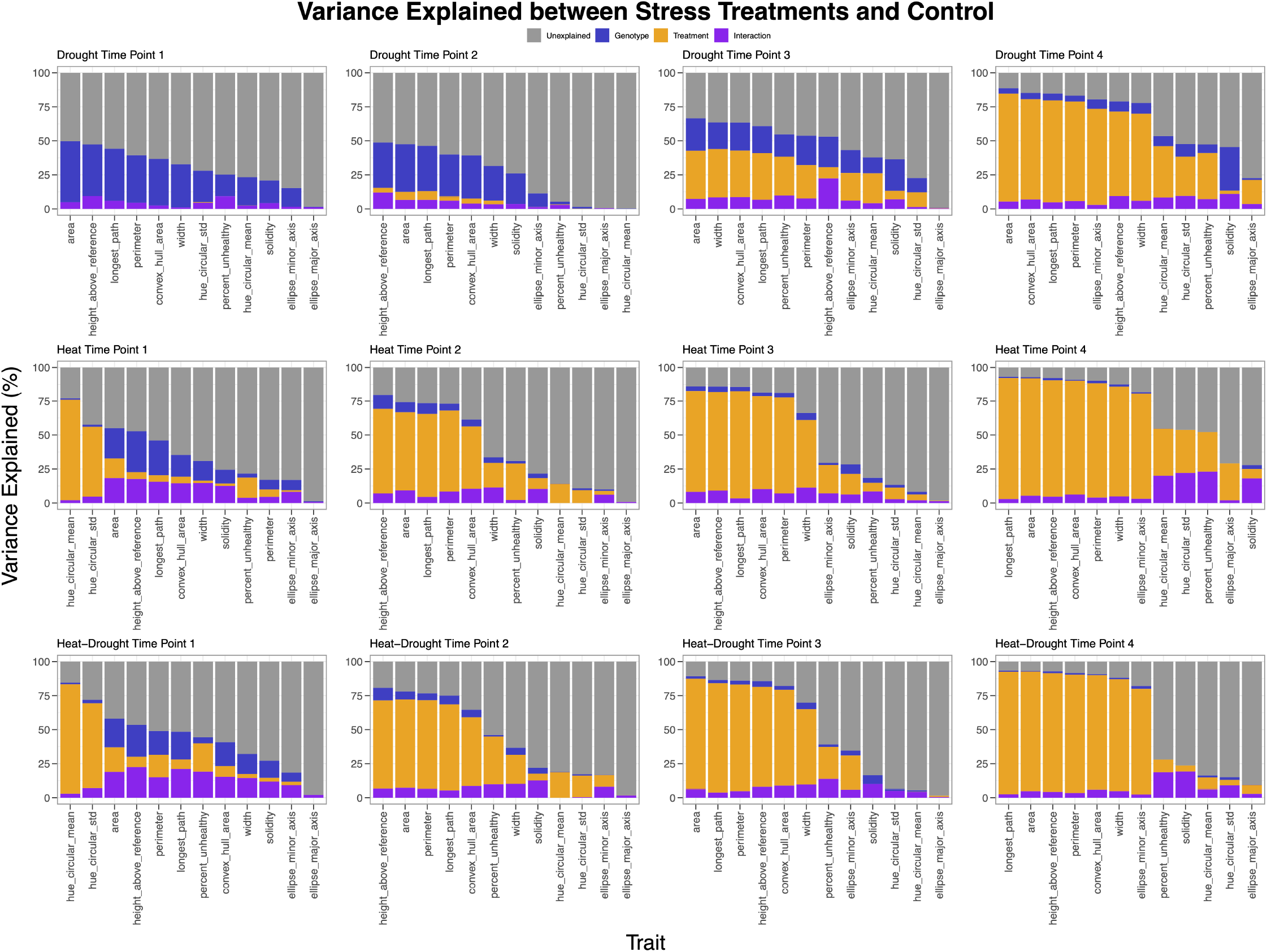
(variance explained): Stacked bar graphs of the variance explained between all stress treatments compared to control conditions. The x-axes show different phenotypic measurements output from PlantCV, and the y-axes show the percent variance explained. Gray bars represent unexplained variance, blue bars represent variance explained by genotype, orange bars represent variance explained by treatment, and purple bars represent variance explained by genotype-treatment interaction. **A)** The variance explained between control conditions and the drought treatment. **B)** The variance explained between control conditions and the heat treatment. **C)** The variance explained between control conditions and the combined heat and drought treatment.

### B. distachyon has strong population structure with two main subpopulations separated based on country of origin

Population structure is the presence of a difference in allele frequencies between different groups in the population and it often arises from physical separation between members of a species either because of distance or barriers (Corander et al., 2008). Population structure is important to identify because it could lead to errors in genetic analyses like GWAS if a homogenous distribution of alleles throughout the population is assumed (Corander et al., 2008).

Using SNPs identified from the GBS data, a PCA was done to examine the population structure of the *B. distachyon* accessions in this study (Figure 8). PCA shows two main clusters (with a few accessions not in either cluster), which indicates that there is strong population structure present in this population of *B. distachyon* accessions (Figure 8). These results support previous conclusions that found strong population structure in *B. distachyon* populations (Draper et al., 2001; Garvin et al., 2008; Filiz et al., 2009; Tyler et al., 2016). Clustering is almost exclusively by country of origin, with the accessions from Spain and Turkey making up the two main subpopulations (Figure 8). Fixation index (Fst) calculations between the two main subpopulations shown in the PCA also confirmed genetic separation between these subpopulations (Weir and Cockerham mean Fst estimate = 0.4014).

**Figure 8.**
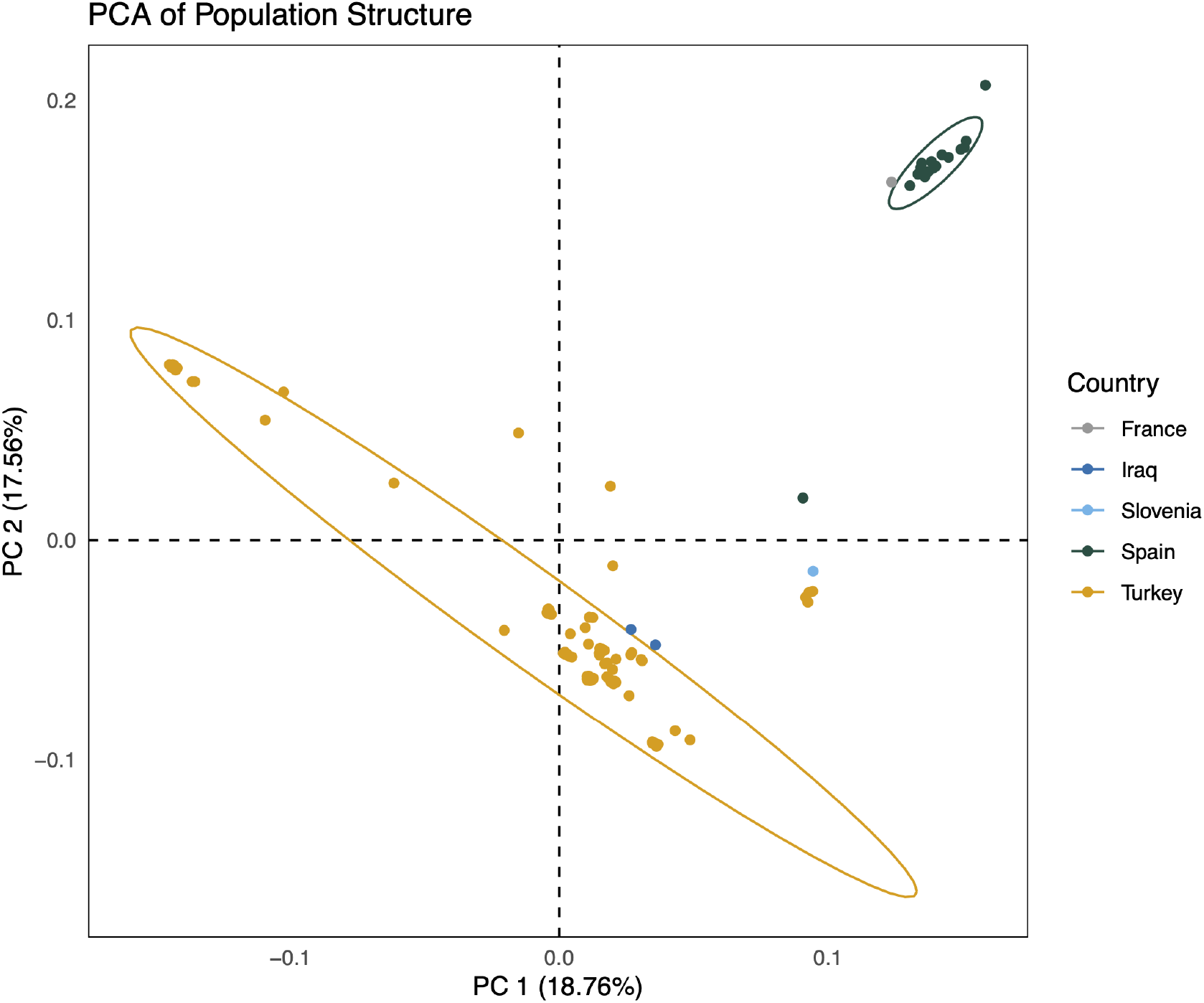
(population structure PCA): Principal component analysis (PCA) of *B. distachyon* population structure. This PCA incorporates SNP information, called using Stacks, for 147 *B. distachyon* accessions. Graph shows principal component 1 (PC 1), which explains 18.76% of the variance, on the x-axis, and principal component 2 (PC 2), which explains 17.56% of the variance, on the y-axis, with 95% confidence ellipses included. Points are colored by the country of origin for each accession.

### B. distachyon population climate of origin is associated with height and plant tissue damage in heat and drought stresses

To see if there was correlation between more ‘stress tolerant’ accessions and the climates that they were collected in, the correlation between the climate at each *B. distachyon* accession’s collection location (climate of origin) and its phenotypic response under stress was examined. Although the *B. distachyon* accessions assessed in this study are generally located in hot, arid regions of the world, there is still considerable variation in climate of origin. Given the hypothesis that plants have adapted to the environment in which they have evolved in (Anderson et al., 2011; Mitchell-Olds et al., 2007), a goal of this research was to determine if there was correlation between phenotype and climate of origin that might suggest that adaptations to different climates could be driving genetic and phenotypic differences in *B. distachyon*. Using data obtained from WorldClim (Fick and Hijmans, 2017), 19 bioclimatic variables plus elevation data from each accession’s collection location were included in this analysis. A correlation plot of the bioclimatic variables shows strong correlations between different groups of variables, which help to generally describe the climates of origin for the *B. distachyon* accessions (Figure 9). The correlations between the bioclimatic variables indicate that the accessions come from climates that are generally hot and dry, or cool and wet at different times of the year. The strong positive relationships between Precipitation of Driest Month (bio14), Precipitation of Driest Quarter (bio17), and Precipitation of Warmest Quarter (bio18), as well as the strong negative relationship between Mean Temperature of Warmest Quarter (bio10) with the three previously mentioned precipitation-related variables (bio14, bio17, bio18) indicate that the warmest time of year in the climates of origin of this *B. distachyon* population coincides with the driest time of year (Figure 9). The strong positive correlation between Precipitation of Wettest Quarter (bio16) and Precipitation of Coldest Quarter (bio19) demonstrates that the wettest time of the year is also the coolest time of the year (Figure 9). Also, higher elevations are associated with cooler temperatures in the *B. distachyon* climates of origin (as is generally expected), as indicated by the negative relationship between Elevation and Annual Mean Temperature (bio1), Max Temperature of Warmest Month (bio5), Min Temperature of Coldest Month (bio6), Mean Temperature of Warmest Quarter (bio10), and Mean Temperature of Coldest Quarter (bio11) (Figure 9).

**Figure 9.**
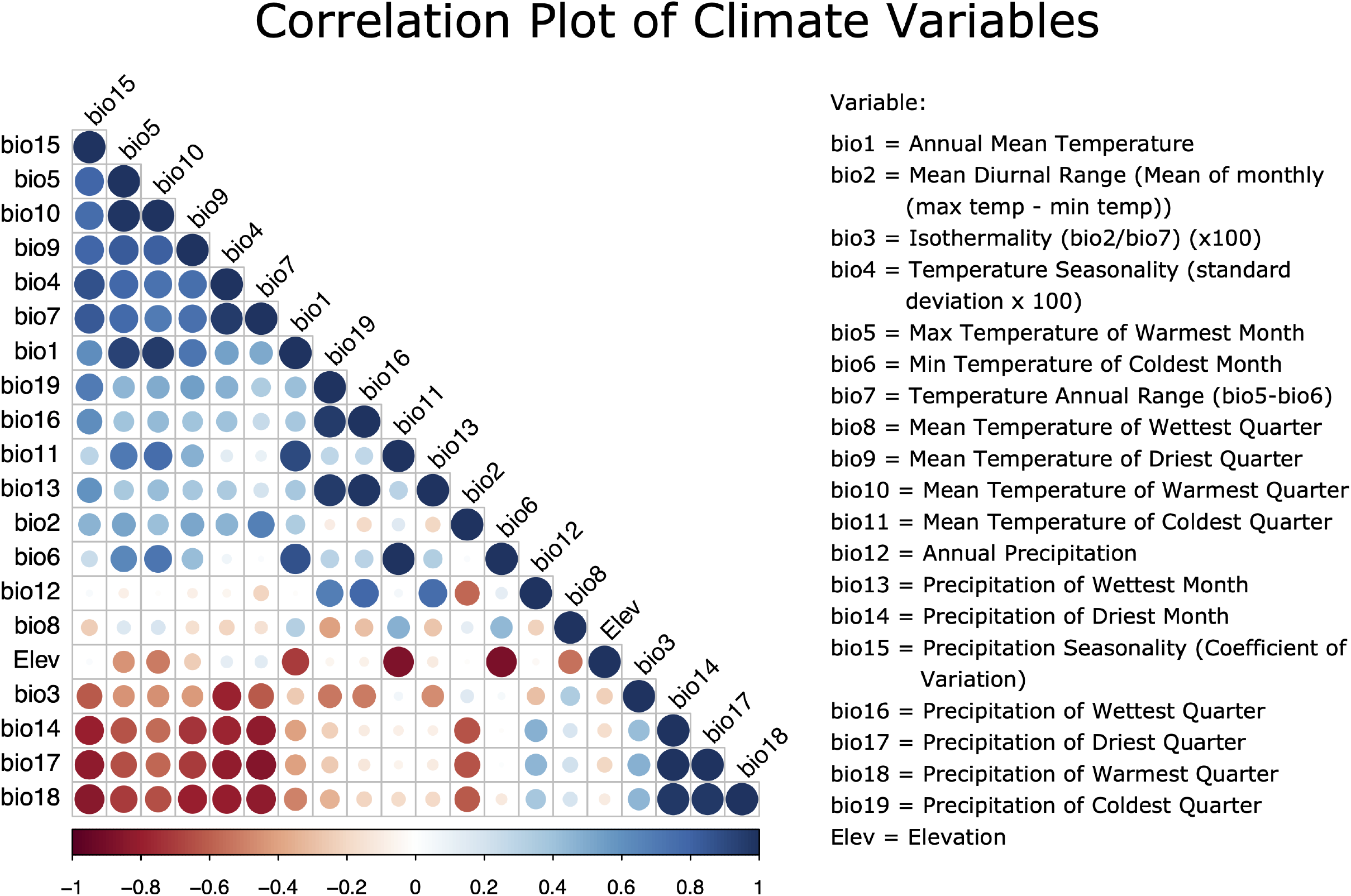
(climate correlation plot): The correlation between variables used in the PCA of climates of origin. Larger, darker circles represent stronger correlation between variables and smaller, lighter circles represent weaker correlations between variables. The plot is ordered by “first principal component order” (FPC) by the corrplot package in R (Wei and Simko, 2017).

A principal component analysis (PCA) of climate data (19 bioclimatic variables) showed that climatic differences grouped by collection country of origin, similar to the population structure clusters (Figures 8 and 10). The PCA indicated that the clear differences in climate of origin across the *B. distachyon* population explain a large portion of the variation across the climate variables (~70%; Figure 10). There were two main climate clusters, one mostly representing the *B. distachyon* accessions collected in Turkey (dark blue points; Figure 10) and the second clustering of the data points predominantly representing the accessions collected in Spain (green points; Figure 10). This separation suggests meaningful differences in the climates of these two native growth regions for *B. distachyon*. Over time, these climatic differences could contribute to selective pressures that drive differences in abiotic stress tolerance between *B. distachyon* accessions. Since there is only one accession from France and Slovenia, and two from Iraq in the 147 *B. distachyon* accessions with climate data available, there are not enough data to make definite conclusions about the climate of these regions compared to the others (Figure 10).

**Figure 10.**
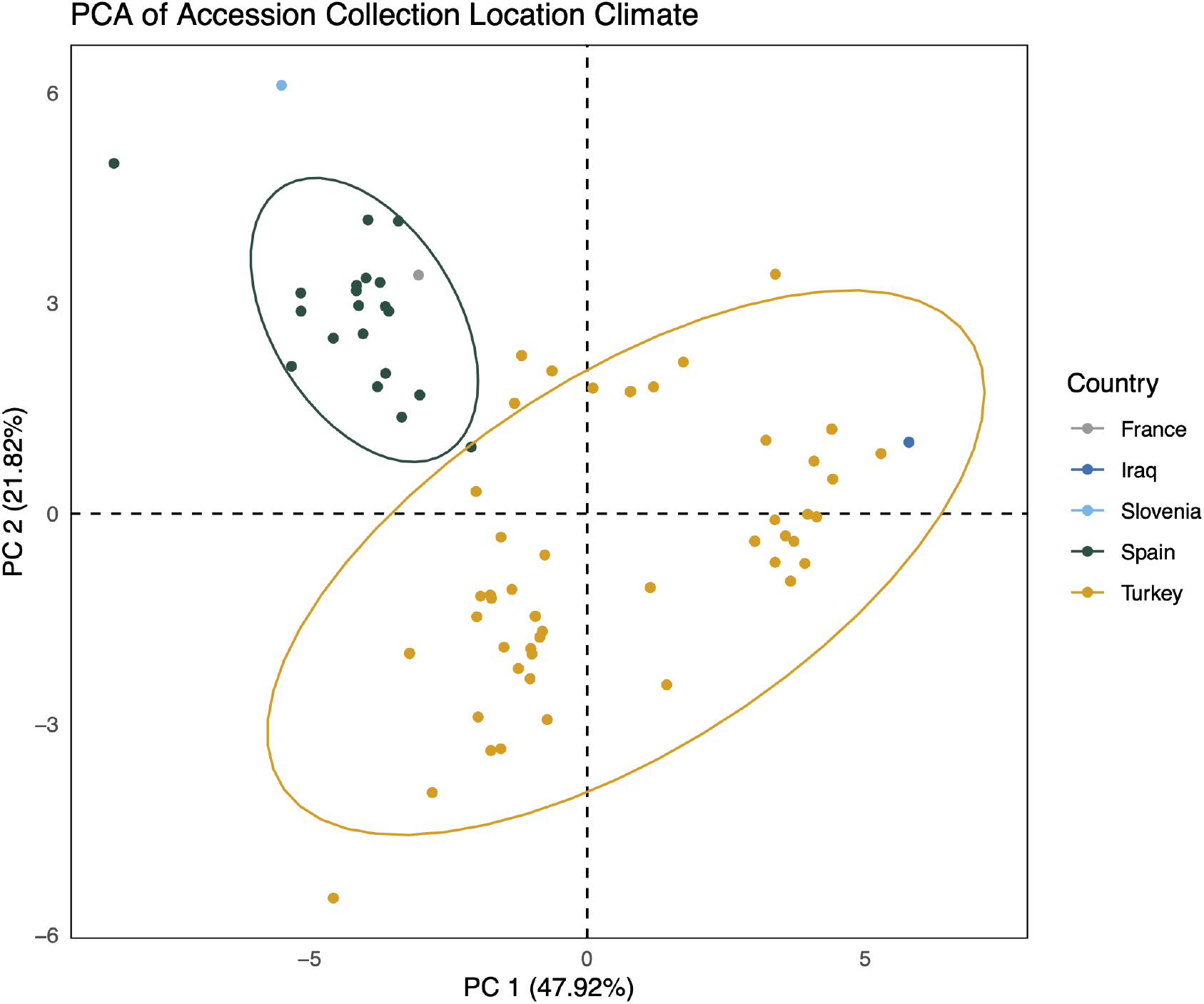
(climate PCA): Principal component analysis (PCA) of *B. distachyon* collection location climates, using data downloaded from WorldClim. This PCA incorporates 19 bioclimatic variables describing in detail the climate in the location where the 147 of the 149 *B. distachyon* accessions were collected (specific collection location is unavailable for two accessions). The graph shows principal component 1 (PC 1), representing 47.92% of the variation explained, on the x-axis and principal component 2 (PC 2), representing 21.82% of the variance explained, on the y-axis, with 95% confidence ellipses included.

Further evaluation of the WorldClim data revealed significant associations between climate of origin and *B. distachyon* phenotypes under abiotic stress conditions (p-values < 0.05, Spearman correlation test). The first three climate principal components (PCs) were assessed because they each explain over 15% of the variance in accession climate of origin and a total of 89.08% of the variance explained (Supplemental Figure 5). All other PCs individually explain less than 5% of the variance (Supplemental Figure 5), which means they likely offer less than one variable’s worth of information and were therefore excluded from further analysis (Swan et al., 1995; Legendre and Legendre, 2012). There were no significant associations between climate PC 1 and any phenotypes, however, climate PC 2 is significantly associated with percent damage in the heat-drought combination (p = 0.00191), and climate PC 3 is significantly correlated with percent damage in the heat treatment (p = 0.03258), as well as height in all three stress treatments (drought, p = 0.01328; heat, p = 0.01118; heat-drought p = 0.01657). The variable contributions to each of these PCs (Supplemental Figure 6), were examined to understand which of the 19 bioclimatic variables contribute to which PCs. For this PCA, loadings greater than the cutoff of ±0. 2236 are considered significant (Swan et al., 1995; Legendre and Legendre, 2012). The variables that contribute most strongly to climate PC 2 are Minimum Temperature of Coldest Month (bio6), Elevation, and Mean Temperature of Coldest Quarter (bio11), and the variables that contribute most strongly to PC 3 are Annual Precipitation (bio12), Precipitation of Wettest Quarter (bio16), and Precipitation of Wettest Month (bio13) (Supplemental Figure 6). Both PC 2 and PC 3 have variables of all types (temperature, precipitation, combination, elevation) that contribute significantly to these principal components. However, the variables contributing significantly to PC 2 seem to be broadly related to temperatures in the coldest month/quarter of the year and precipitation in the driest month/quarter of the year, while the variables contributing significantly to PC 3 seem to be broadly related to temperatures in the wettest month/quarter of the year, precipitation in the wettest month/quarter of the year, and temperature seasonality (Supplemental Figure 6). These associations overall indicate that there are significant correlations between climate of origin and *B. distachyon* phenotype in abiotic stress conditions, which could indicate local adaptation that is driving the diverse stress responses of these accessions.

### Genome wide association mapping found SNPs significantly associated with height, area, and percent tissue damage under stress conditions

A genome wide association study was conducted to identify genetic loci associated with changes in plant phenotypes under heat and drought treatments. To assess the genetic basis of heat and drought responses in *B. distachyon* over time, separate GWA analyses were done using data from each of the four Time Points previously mentioned (Time Point 1 on Imaging Day 4, Time Point 2 on Imaging Day 7, Time Point 3 on Imaging Day 10, and Time Point 4 on Imaging Day 15; the final day with data for all treatments). A separate GWAS was done for each stress treatment by taking the average of measurements for the specific trait of interest across all replicates for a given treatment at that time point, and taking the difference of those averages compared to the average of the replicates under control conditions, and using this difference as the trait measurement for the GWAS. This was done for three phenotypic measurements: plant height, plant shoot area, and percent damage.

In total, 16 SNPs were significantly associated (p < 1.71e-5) with three traits: height, area, and percent damage (Figure 11, Supplemental Figure 7, Supplemental Figure 8). The average linkage disequilibrium (LD) decay was found to be 100,000 bp at an r^2^ = 0.2 (Supplemental Figure 9), so candidate genes could be found within this range, upstream or downstream from a significant SNP. However, such a range can often contain thousands of genes, and since this was a preliminary exploratory analysis of these genes, in this study only the two adjacent genes closest to a significant SNP were assessed.

**Figure 11.**
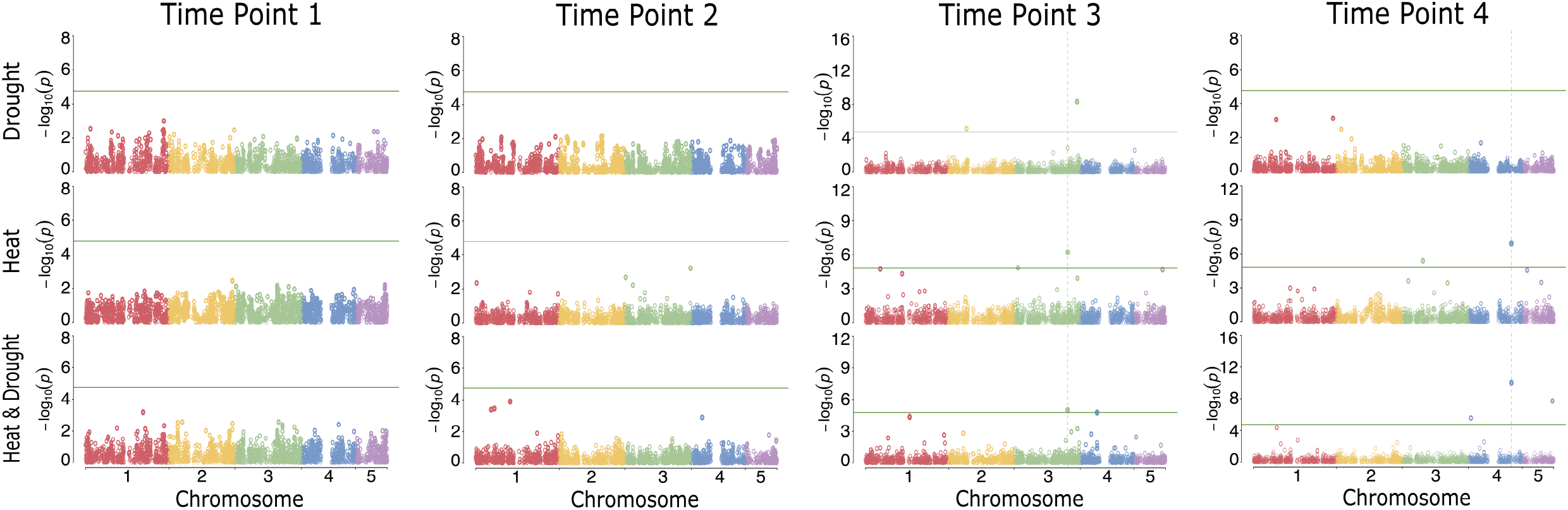
(height Manhattan plots): Manhattan plot of GWAS discovery results for height. Each point represents a SNP. The height of the SNP on the y-axis represents the strength of association with plant height, expressed as −log10(p-value). Points are colored by chromosome, and appear darker and more solid with lower p-values (stronger association with plant height). The green horizontal line represents the significance cutoff and is calculated by dividing 0.05 by the number of SNPs included in the GWAS: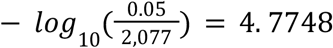

Eight SNPs were significantly associated with height; four at Time Point 3 and four at Time Point 4 (Figure 11). At Time Point 3, there were SNPs significantly associated with height in all three stress treatments, with one SNP overlapping between the heat and heat-drought treatments (Figure 11). At Time Point 4, SNPs were significantly associated with height in the heat and heat-drought treatments, with one SNP found to be significant in both (Figure 11). Five SNPs were significantly associated with area, with one significant SNP found at Time Point 2 in the heat-drought treatment, and four at Time Point 4 in the heat and heat-drought treatments (Supplemental Figure 7A). Three SNPs were significantly associated with percent damage, and all were at Time Point 3 in the drought treatment (Supplemental Figure 7B).

A table of genes closest to SNPs is available in Supplemental Figure 8. These genes closest to significant SNPs are discussed further in the Discussion section and are potential targets to alter plant growth and architecture under abiotic stress conditions.

## Discussion and Conclusions

Current elite crops have been bred to be extremely high-performing but not necessarily very tolerant to fluctuations in climate or weather patterns that expose crops to abiotic stresses (Mickelbart et al., 2015). Accordingly, it is valuable to explore the natural variation of phenotypes in weedy relatives to these major crops, to exploit their diversity in stress tolerance and improve that of current elite crops (Mickelbart et al., 2015). *B. distachyon* is an example of a weedy relative of major food and biofuel crops with immense natural phenotypic variation (International Brachypodium Initiative, 2010). Studying this powerful model C_3_ grass to assess its natural variation and understand genetic loci associated with its abiotic stress responses could aid in improving abiotic stress tolerance in the elite crops that feed and fuel Earth’s growing population. Also, the anticipated increase in drought and heat stress as a result of climate change and the distinct impact of stresses in combination underscores the importance of phenotyping plants under multiple stresses that frequently co-occur.

The differences in responses to individual versus combined abiotic stresses is central to this study. For plant biomass accumulation (plant shoot area and height), the combination of heat and drought stress is generally the most detrimental for *B. distachyon* (Figure 6, Supplemental Figure 3). *B. distachyon* accessions accumulated less biomass in the combination of stress conditions compared to when exposed to heat or drought stresses individually (Figure 6, Supplemental Figure 3). Plants in heat stress were also much smaller than in drought stress and closer in size to plants in heat and drought, but on average were still able to grow larger than in the combination stress treatment (Figure 6, Supplemental Figure 3). These results alone might indicate that the combination of heat and drought stresses is more severe than individual stresses, but when considering all results of this study, particularly tissue damage, heat stress seems to be more detrimental than the severe drought treatment applied here or the combination of these two stresses for *B. distachyon*. Further, when comparing all measured phenotypes in this study, the drought stress is distinct from both heat-related stresses, while overlap in responses was observed between the heat and heat-drought stresses (Figure 3). Variance in phenotype explained by genotype, treatment, and the genotype-treatment interaction show the treatment effect appearing earlier for the heat and heat-drought treatments compared to the drought treatment (Figure 7), suggesting that heat is more damaging than drought in this study. The treatment effect is also much larger in both heat-related treatments than in the drought treatment (generally 7-12 percent larger) (Figure 7).

Fitting with the variance explained data, plant tissue damage in *B. distachyon* was generally the highest in response to heat stress in comparison to either drought or the combination of heat and drought (Figure 4). Therefore, although past studies suggest that the responses to the combination of drought and heat stress would generally be more severe than individual stresses (Rizhsky et al., 2004, 2002; Rasmussen et al., 2013; Choudhury et al., 2017; Mittler, 2006; Balfagón et al., 2020; Atkinson et al., 2013), our data show that for the plant tissue damage trait, heat stress is the most severe stress for *B. distachyon* under the tested conditions. These results suggest that *B. distachyon* may have stress avoidance strategies to survive hot and dry conditions by accumulating less biomass under drought stress and the combined heat and drought stress, while still maintaining healthy tissue capable of photosynthesizing. Conversely, *B. distachyon* does not appear to be well equipped to survive under heat stress alone, and therefore, although it is able to accumulate marginally more biomass in heat than in the combination of heat and drought, this plant tissue is extremely stressed.

As previously mentioned, abiotic stresses like heat and drought are often found to occur simultaneously in nature, so it is possible that *B. distachyon* has adapted to maintain photosynthetically active tissue under a combination of drought and heat but not under heat alone. In fact, the native growth region of *B. distachyon* is in the Mediterranean and Middle East, regions of the world that are generally warm or hot and very arid (Draper et al., 2001; Garvin et al., 2008; Filiz et al., 2009). According to the most recent Köppen-Geiger climate classification, the collection locations of the *B. distachyon* accessions examined in this study are generally classified as arid or temperate, with dry summers (Peel et al., 2007). This means that this region is warm and typically does not receive much precipitation, so *B. distachyon* would likely very rarely be exposed to hot but well-watered conditions, similar to the heat stress alone treatment, in its climates of origin. This is corroborated by the climate data presented from WorldClim, which shows that the climates of origin of these *B. distachyon* accessions are hot and dry, or cool and wet in different seasons (Figure 9), and overall receive relatively low amounts of precipitation annually. The times of the year with the most precipitation and the warmest temperatures do not coincide, supporting the conclusion that these *B. distachyon* accessions are likely not adapted to hot and wet conditions. Consequently, it may be that under heat stress alone, *B. distachyon* is primed to encounter drought as well, and is more susceptible to osmotic stress when well-watered. Conversely, it could also be that when *B. distachyon* is exposed to drought, as it often is in its native climates, it is also primed for heat stress, so when well-watered it is not primed to maintain healthy tissue in heat. This is an important avenue to explore in future studies to help elucidate the mechanisms by which *B. distachyon* responds to different stresses and their combinations.

While the climates of origin of these *B. distachyon* accessions have similarities – generally dry and warm or hot summers with cooler, wetter winters – the climate data from WorldClim presented also show variation in these climates of origin (Figure 10). Analyses of these climate data and *B. distachyon* phenotype under drought and heat stresses revealed significant associations between the climate of origin and accessions’ responses to these stresses. This suggests that these accessions may be locally adapted to their climate of origin, which is driving their responses to these heat and drought stresses. Two accessions, BdTR10E and BdTR5J, illustrate this idea clearly. Both accessions are from Turkey, but from different geographic regions with distinct climates, and contrasting responses to the drought and heat stresses in this study. BdTR10E was one of the two accessions identified as being most tolerant to all three stress treatments, compared to BdTR5J, which was identified as being one of the two *least* tolerant to these stresses (Supplemental Figure 2). BdTR10E is from the Southeastern Anatolia Region of Turkey, East of the city of Gaziantep (Filiz et al., 2009), which is a relatively low-lying area of the country (Figure 1, Supplemental Figure 1). The collection location of BdTR10E sits at an elevation of 448 m above sea level and has a hot and dry climate with low amounts of precipitation (Supplemental Figure 1). In contrast, BdTR5J is from the Central Anatolia Region of Turkey, North of the city of Ankara, in the Köroğlu Mountains (Figure 1, Supplemental Figure 1) (Filiz et al., 2009). It was collected at an elevation of 1556 m above sea level, in a climate that is not as hot overall, is not as hot during the dry season specifically, and receives more precipitation even during the warmer season (Supplemental Figure 1). These data show clear differences between the climates of origin of BdTR5J and BdTR10E, specifically that BdTR10E was collected in an area with a much hotter and drier climate than where BdTR5J originates from. This is especially interesting considering that BdTR10E performed well under drought and heat stress conditions, and BdTR5J performed poorly under these stress treatments. This could hint that these accessions are locally adapted to their climates of origin. However, reciprocal transplant experiments would be required to test this hypothesis to see if there are higher rates of survival and fitness for *B. distachyon* accessions in their home environments.

This study found a number of genetic loci significantly associated with height, shoot area, and the percent damage in the different abiotic stress treatments compared to control conditions (Figure 11, Supplemental Figure 7). These results could point to genes that regulate *B. distachyon”*s responses to drought, heat, and the combination of drought and heat. While two SNPs were found to be significantly associated with height in both the heat treatment and the heat-drought in combination, most significant SNPs were distinct between the different stress treatments, which suggests that the combined stress may activate a unique set of stress pathways compared to each stress individually (Figure 11, Supplemental Figure 7). This suggests that different genes are associated with the responses to the tested abiotic stresses. Since heat and drought often activate different mechanisms by which plants respond to stresses (Atkinson and Urwin, 2012; Rizhsky et al., 2004, 2002; Mittler, 2006), it would not be surprising to discover that different genes are involved in drought tolerance compared to heat tolerance, as well as the combination of these two stresses. Eight SNPs were significantly associated with height at different times during the experiment (Figure 11). Notable candidate genes located near these SNPs significantly associated with height that are known to be involved in plant abiotic stress responses include a sugar transporter (Kaur et al., 2021; Saddhe et al., 2021; Gautam et al., 2019; Gupta and Kaur, 2005), a Replication Factor-A protein (Danilevskaya et al., 2019), an autophagy (ATG) protein (Zhou et al., 2014; Bassham et al., 2006), and a fasciclin 1 (FAS1) domain (Seifert, 2018) (Supplemental Figure 8). A particularly interesting candidate gene in this case is the FAS1 domain, which is in close proximity to the SNP significantly associated with height in both the heat and heat-drought treatments (Figure 11). Fasciclin domains are a cell adhesion domain that is found in insects, animals, bacteria, fungi, algae, and is found as a large family of fasciclin-like arabinogalactan proteins (FLAs) in higher plants (MacMillan et al., 2010; Faik et al., 2006). Some FLAs have been linked to secondary cell wall synthesis in Arabidopsis stems, and with wood formation in tree trunks and branches, which suggests a role in the development of plant stems, and research into this gene family has found that FAS domain FLAs contribute to stem strength in plants by regulating cellulose deposition and affecting the integrity of the cell-wall matrix (MacMillan et al., 2010). Studies have shown that FLA genes are involved in plant abiotic stress response, including findings showing that some FLA genes were up-or down-regulated in response to abiotic stress including heat and dehydration in wheat (*Triticum aestivum*) (Faik et al., 2006), and that rice (*Oryza sativa L.*)FLA genes also displayed differential expression patterns induced by abiotic stresses (Ma and Zhao, 2010). In addition to abiotic stress-related FLA genes having been identified in close genetic relatives of *B. distachyon*, a study examining FLA transcript levels in response to temperature stress in *B. distachyon* found that multiple FLA genes were upregulated at high temperature compared to the control (Pinski et al., 2019). This candidate gene is especially noteworthy considering the role that FLA genes play in cell wall development and stem strength in plants and that this SNP is significantly associated with height.

Five SNPs were significantly associated with shoot area throughout the experiment (Supplemental Figure 7A). Potentially interesting candidate genes known to be involved in abiotic stress responses in plants include a Syntaxin 7 (STX7) t-SNARE domain containing protein (Zhu et al., 2002; Kwon et al., 2020; Chen et al., 2019), an Inositol 5-Phosphate gene (Perera et al., 2008; Burnette et al., 2003; Na and Metzger, 2020; Jia et al., 2019), and a Calmodulin-binding Transcription Activator (CAMTA) (Galon et al., 2010; Pandey et al., 2013; Noman et al., 2021; Yang et al., 2010; Shen et al., 2015). One SNP, found on Chromosome 3 and significantly associated with shoot area in the heat treatment, is located within a gene that codes for a CAMTA transcription factor (TF) (Supplemental Figure 7A). CAMTAs are a vital family of proteins that have an impact on diverse cellular processes by regulating gene expression (Pandey et al., 2013) either by binding to DNA directly or by interacting with other TFs (Finkler et al., 2007). CAMTAs have been linked to multiple different signaling pathways, including the Ca^2+^-signaling pathway, which plays a crucial role in stress signaling and adaptation in response to a variety of stresses, including abiotic stress (Finkler et al., 2007; Yang and Poovaiah, 2000; Noman et al., 2021). Studies have shown that mutations in CAMTA genes have greatly impacted responses to a wide variety of abiotic stresses, including drought and heat in *Arabidopsis thaliana* and *Oryza sativa*, among others (Pandey et al., 2013; Noman et al., 2021; Shen et al., 2015; Finkler et al., 2007; Galon et al., 2010). This indicates that CAMTA genes are a potential target to alter plant growth in response to heat, drought, and other abiotic stresses.

Three SNPs were significantly associated with percent shoot damage under drought conditions (Supplemental Figure 7B). One notable candidate gene located near the SNPs significantly associated with percent damage is RAS-related protein RABE1A (Supplemental Figure 8). Rab proteins are guanosine triphosphate (GTP)ases that regulate vesicle trafficking (Chen et al., 2021) by cycling between two forms: an active, membrane-bound GTP, and an inactive, cytosolic GDP (guanosine diphosphate) (Tripathy et al., 2021). Rab genes are subdivided into eight clades, and RabE proteins (also referred to as Rab8) are involved in transport between the Golgi apparatus and the plasma membrane (Speth et al., 2009; Zheng et al., 2005; Mayers et al., 2017; Chen et al., 2021). Studies have found that RabE/Rab8 proteins are involved in plant development and autophagy, as well as abiotic stress responses (Tripathy et al., 2021; Chen et al., 2021; Zhang et al., 2018). In *A. thaliana*, for example, RabE1c expression is highly induced by drought stress, and overexpression of RabE1c conferred increased drought tolerance in *A. thaliana* and chinese cabbage (*Brassica rapa* ssp. *pekinensis’*) (Chen et al., 2021). This points to RabE/Rab8 genes as being potential targets for genetic modification to improve plant drought tolerance.

This dataset can be leveraged for future studies into uncovering the mechanism of heat and drought responses in *B. distachyon*.

## Materials and Methods

### Plant Material

Diploid accessions of *B. distachyon* from the USDA and the Mur Lab (University of Aberystwyth) were used in both phenotyping experiments. The population of *B. distachyon*accessions used in this project was collected throughout this model plant’s native range in the Mediterranean and the Middle East (Draper et al., 2001; Garvin et al., 2008; Filiz et al., 2009; Tyler et al., 2016). The majority of accessions were collected in Turkey and Spain, but the distribution of this population’s collection locations can be seen in Figure 1. More detailed information regarding the collection locations of these accessions can be found in the Supplemental Figure 1. 137 accessions were included in the experiment with drought and control conditions, and 144 accessions were used in the experiment with heat and heat-drought conditions. These were overlapping sets of *B. distachyon* accessions, with 132 being included in both experiments. Approximately 4 replicate plants of each accession were planted for every treatment.

### Plant Growth and Application of Stress Treatments

The *B. distachyon* accessions in each experiment were split into two groups and planted in plug trays on either day 1 or day 2 of both experiments (control and drought; heat and heat-drought). Splitting the accessions into two planting groups made it possible to compare plants of the same age in images taken throughout the experiment, since plants were imaged every other day. After planting, accessions were kept in a Conviron growth chamber for 10 days to germinate, before being transplanted into prefilled 4-inch pots that were labeled with barcodes encoding accession identification, water treatment group and a unique pot identification number. The repotted plants were then returned to their original Conviron growth chamber for two more days before being loaded into the Bellwether Phenotyping Platform’s Conviron growth chamber at 12 days after planting (DAP).

In both experiments, plants were grown under 14h-photoperiod (14h day/10h night) with a light intensity of 200 μmol/m^2^/s and relative humidity at 50%. In the control and drought experiment, the temperature was 22°C during the day and 18°C at night, and in the heat and heat-drought experiment it was 35°C during the day and 30°C at night. Water treatment was done by watering to 100% of the target weight for the well-watered treatment, and 20% of the target weight for the water-limited (drought) treatment. For the first four days on the phenotyping platform, all plants (regardless of treatment) in both experiments were watered to 100%, with the 20% water-limited treatment being imposed on the plants in the drought treatment group on the fifth day. Target weights for watering were calculated using the method described in Fahlgren et al. (Fahlgren et al., 2015).

### Image Processing and Extraction of Trait Information

RGB (red, green, blue) images of all individual plants were captured every other day in the Bellwether Phenotyping Facility at the Donald Danforth Plant Science Center, acquiring side-view images from four angular rotations (0°, 90°, 180°, 270°). To ensure the entire plant was included in each image, optical zoom was adjusted throughout the experiment as the plants grew in size.

Images were analyzed using the open-source image analysis software PlantCV (Gehan et al., 2017). PlantCV v3.8 (Fahlgren et al., 2020) was used to analyze each camera angle and zoom level. To segment the plant from background, the RGB image was converted to the HSV and LAB color-spaces and the saturation and blue-yellow channels were isolated. Thresholds were applied to the saturation channel and the blue yellow channels and the two binary images were combined. The fill function of PlantCV was used to fill in any noise (objects smaller than 50 pixels) that were left over after combining the two binary images. The cleaned binary image was used to mask the original RGB image so that the pot, carrier and background were removed. A Region of Interest (ROI) was set around the plant to further isolate it from the background, and only objects that overlapped the ROI were kept for analysis. After segmentation of the target object (plant) from the background, morphological characteristics were extracted using the PlantCV ‘analyze_object’ function (Gehan et al., 2017). Additionally, the PlantCV naive Bayes classifier module (Abbasi and Fahlgren, 2017; Gehan et al., 2017) was trained to label plant pixels as either “healthy” or “unhealthy” to determine what percentage of the plant in each image was stressed by its growth conditions. Training was done by choosing a representative set of 10 images and using the Pixel Inspection Tool in ImageJ (Abràmoff et al., 2004) to gather color value training data for over 1,000 pixels from each of three categories: healthy plant tissue, unhealthy plant tissue, and background (discarded from further analysis). The pixel color data was then used in PlantCV to train the naive Bayes classifier, which was included in the Python scripts run over each image to assign the plant pixels to the appropriate category of interest. The percentage of damaged tissue was calculated by dividing the shoot area classified as unhealthy by the total shoot area. Representative images of a few of the outputs from PlantCV, including the naive Bayes classifier, used in the analysis of this data are shown in Figure 12.

**Figure 12.** (plantcv output examples): Representative input and output images from PlantCV. **A)**Example of a side-view image captured during the experiments that would be used as an input for the PlantCV image-analysis pipeline. **B)**Example of the “height_above_reference” output from PlantCV. **C)**Example of the “area” output from PlantCV **D/E)**Example of the naive Bayes classifier categorizing plant pixels as either healthy (blue) in panel **D** or unhealthy (orange) in panel **E**.

To correct for the different zoom levels in the images, scaling factors for both height and area were calculated in R (R Core Team, 2020) using a reference object of known size as was done in Fahlgren et al. (Fahlgren et al., 2015). The dimensions of the known object were measured in centimeters and it was imaged with the same imaging setup and at the same zoom levels used in the *B. distachyon* phenotyping experiments. Height and area of the known object were then measured in pixels, which allowed a conversion factor to be calculated. This was used to convert pixel measurements from PlantCV to centimeters or centimeters squared, which allowed pixel heights and areas that represent plant phenotypes to be compared across zoom levels in downstream analysis. Due to the time it takes to image the plants on the phenotyping platform, only half of the plants were imaged on each day, but plant growth was staggered to account for this. Half of the plants were assigned to be imaged on odd days, and the other half on even days. For analysis purposes, data was labeled by ‘Imaging Day’. For example, days 1 and 2 of the experiment are Imaging Day 1, days 3 and 4 Imaging Day 2, and so on.

R analysis scripts and Python scripts for all image analysis and trait extraction done with PlantCV are available in a public GitHub repository at this link: https://github.com/danforthcenter/brachypodium-heat-drought-paper.

### High Resolution Spatial Climate Data

The climate data used for the climate analysis in this study were obtained from WorldClim V.2.1, a long-term global climate dataset (Fick and Hijmans, 2017). WorldClim provides monthly high-resolution spatial gridded data of 19 bioclimatic variables for a 30 year reference period between 1970 and 2000 at a resolution of 30 arcseconds, which is approximately 1 km^2^ on the Earth’s surface (Fick and Hijmans, 2017). These data are interpolated using in situ observations at hydro-meteorological stations from various programs, data sources and entities, including: the global historical Climatology Network (ghCN), the World Meteorological Organization (WMO) and the International Center for Tropical agriculture (CIaT), among others (Fick and Hijmans, 2017). The euclidean distance between each accession’s collection location and the WorldClim data points were calculated to find the closest available climate data for each accession’s climate of origin. Climate data from WorldClim was adequately close (euclidean distance <√2km) to 147 out of the total 149 accessions examined in this study (Supplemental Figure 1).

The climate data was used to conduct a PCA in R, and the loadings (the amount that each variable contributes to a principal component) were examined. Since the sum of squares of all loadings for an individual PC must sum to one, the formula 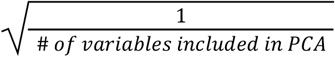 can be used to find the cutoff for determining a significant loading for a PC because this value represents what the loadings would be if all variables contributed equally to that principal component (Swan et al., 1995; Legendre and Legendre, 2012). Any variables with a loading greater than this cutoff value contribute more than one variable’s worth of information and can therefore be treated as an important contributor to that PC. In this case, the cutoff for significant loadings is |0. 2236|.

### Low-Coverage Sequencing

Plant tissue was collected from each accession following the completion of the phenotyping experiments. Genomic DNA was extracted and quantified, and genotype-by-sequencing (GBS) libraries were constructed using the protocol described in Huang et al. (Huang et al., 2014). These libraries were sequenced by the Genome Technology Access Center at Washington University in St. Louis. GBS data for this population of *B. distachyon* is available at the Short Read Archive (SRA) PRJNA312869.

### SNP Calling and Population Structure Estimation

To call SNPs (single-nucleotide polymorphisms) from the GBS data collected in this experiment, the Burrows-Wheeler Aligner (BWA) was first used to align all accession genomes to the *B. distachyon* reference genome (Li, 2013). BWA version 0.7.12-r1039 was used (Li, 2013). The most commonly used reference accession for *B. distachyon* is Bd21-0, and the newest genome assembly and annotation for this accession (version 3.2 from 2020) that can be found on Phytozome was used for alignment (Goodstein et al., 2012; Haas et al., 2003; Smit et al., 1996-2010; Yeh et al., 2001; Salamov and Solovyev, 2000). Next, the SAMtools version 1.11 view function was used to convert the SAM (Sequence Alignment/Map) files output by BWA to BAM (Binary Alignment Map) files, which are the binary equivalent of a SAM file (Danecek et al., 2021). The BAM files were then sorted using the sort function from SAMtools, which sorts genome alignments by leftmost coordinates (Danecek et al., 2021). These sorted BAM files were then used in ‘gstacks’, a Stacks program (Catchen et al., 2011, 2013), in the reference-based mode, to call SNPs at each locus, relative to the reference genome, and genotype each individual at every SNP identified (Catchen et al., 2011, 2013). Gstacks then phased these SNPs into a set of haplotypes for each individual at every locus (Catchen et al., 2011, 2013). SNPs called by gstacks and a population map were then input into ‘populations’, another Stacks program, to compute population genetics statistics (Catchen et al., 2011, 2013). In ‘populations’, options used included-t set to 10 to use 10 threads,-r set to 0.8 to set the minimum percentage of individuals in the population required to process the locus, --min-maf set to 0.05 to filter SNPs with a minor allele frequency less than 0.05, --write-single-snp to restrict data analysis to only the first SNP per locus, --plink to output genotypes in PLINK format, --structure to output results in Structure format, and --vcf to output SNPs and haplotypes in Variant Call Format (“vcf’) (Catchen et al., 2013, 2011). Stacks version 2.4 was used for both ‘gstacks’ and ‘populations’ (Catchen et al., 2011, 2013). After this, PLINK v1.90b6.21 was used to do a principal component analysis (PCA) on the output from populations, to find the population structure of *B. distachyon* based on samples in this study (Chang et al., 2015; Purcell, 2020a). The --allow-extra-chr option was used in addition to default parameters for the PLINK --pca function, to allow the *B. distachyon* chromosome codes to be read (Chang et al., 2015; Purcell, 2020a).

Based on the clustering in the PCA to estimate population structure, the *B. distachyon* population in this study was divided into two main subpopulations (with some accessions falling outside these two main clusters). Using vcftools, the fixation index (Fst) values between these subpopulations were obtained (Danecek et al., 2011). Fst is a measure of the genetic variance contained in a subpopulation (the S subscript) relative to the total genetic variance (the T subscript) in a population based on Wright’s F-statistics (Wright, 1965). An Fst value of 0 indicates no genetic differentiation between the subpopulations, while an Fst value of 1 indicates complete differentiation (Bird et al., 2017; Holsinger and Weir, 2009). vcftools v0.1.14 was used with the option --weir-fst-pop to calculate Fst estimates based on Weir and Cockerham’s 1984 paper (Weir and Cockerham, 1984; Danecek et al., 2011).

### Linkage Disequilibrium Decay Estimation

The SNP catalog file in Variant Call Format (“vcf’) was first sorted and converted into bed format using Plink v1.90b6.17, and pairwise linkage decay between markers on the same chromosome was calculated (Chang et al., 2015; Purcell, 2020b). Following this, a bash script was used to extract the distances between the SNPs and the corresponding r^2^ values. Using R, SNPs were categorized into bins of 10 kbps, and means were calculated per bins to estimate linkage disequilibrium decay for this population of *B. distachyon*. Scripts available in a public GitHub repository (https://github.com/danforthcenter/brachypodium-heat-drought-paper).

### Subselection of Accessions used in Data Analysis

There were 137 *B. distachyon* accessions included in the experiment with drought and control conditions, and 144 were included in the experiment with heat and heat-drought conditions. Due to differences in germination and seed availability, 132 accessions were overlapping between both experiments, while the total number of *B. distachyon* accessions examined in either experiment is 149. The 132 accessions overlapping between the two experiments were used in analyses requiring direct comparisons between different treatments. For any analyses using the WorldClim climate data, only accessions with climate data available were used, which were 147 out of 149 included in the phenotyping experiments. In all other analyses, including the GWAS, as many accessions that had complete data were included.

### Statistical Analysis on Extracted Trait Information

All statistical analyses were done in R, using R version 4.0.1 (released June 6, 2020) (R Core Team, 2020). Additional packages used include: corrplot (Wei and Simko, 2017), data.table (Dowle and Srinivasan, 2020), dplyr (Wickham et al., 2021), factoextra (Kassambara and Mundt, 2020), FactoMineR (Lê et al., 2008), GAPIT (Wang and Zhang, 2021), ggplot2 (Wickham, 2016), gplots (Warnes et al., 2020), lme4 (Bates et al., 2015), plyr (Wickham, 2011), raster (Hijmans, 2020), readr (Wickham et al., 2022), reshape2 (Wickham, 2007), tidyr (Wickham, 2020), WorldClimTiles (kapitzas, 2020).

A random effects model (or variance components model) was used to calculate the variance explained by genotype, treatment, and the genotype-treatment interaction in plant traits. The model was fitted using the lmer function from the lme4 package in R (Bates et al., 2015). In this model, a type 3 sum of squares was measured for each plant trait. Those terms were normalized to display a percentage of each plant trait’s total variance explained by the design variables and their interactions.

The GWAS in this study was conducted by using the FarmCPU method implemented in GAPIT Version 3 in R (Liu et al., 2016; Wang and Zhang, 2021). Past studies have established that *B. distachyon* has a very strong population structure (Draper et al., 2001; Garvin et al., 2008; Filiz et al., 2009; Tyler et al., 2016), which was also seen in this study, so the FarmCPU method was used to eliminate false positives due to population structure (Liu et al., 2016). This method also has the ability to include additional covariates to account for population structure or other associations that could lead to false positive results (Liu et al., 2016). Therefore, FarmCPU was first run with “Model.selection = TRUE” so that forward model selection using the Bayesian information criterion (BIC) was conducted to determine the appropriate number of covariates included in the GWAS model (Wang and Zhang, 2021). For all phenotype and treatment combinations, the optimal model included zero covariates for both population structure and climate based on the BIC values, so all FarmCPU runs were done without any manually supplied covariates.

The inputs required to run GAPIT include a genotype file and a phenotype file (Wang and Zhang, 2021). In this case, the phenotype file was created using the data output from PlantCV for three traits: plant height, plant shoot area (to approximate biomass), and the percent damage (to approximate plant stress). To create this file, the average of all images taken of each accession on the chosen imaging day for a given trait was taken, and then the difference between the average of the trait in the stress treatment and the average under control conditions was used as the phenotype value for that accession. This removes the genotype effect and ensures that the ensuing results are due to the treatment effect (GxE). A HapMap file was created by using TASSEL v5.0 (Bradbury et al., 2007) to convert the file containing the genotype and SNP information in variant call format (VCF) to HapMap format using a custom HTCondor job file, which can be found in a public GitHub repository (https://github.com/danforthcenter/brachypodium-heat-drought-paper) to input as the genotype file in GAPIT. Since TASSEL requires the VCF file to be sorted by the position column before it can be loaded and used, this sorting was done by using the SortGenotypeFilePlugin in TASSEL v5.0 (Bradbury et al., 2007), also with a custom HTCondor job file available in the GitHub repository (https://github.com/danforthcenter/brachypodium-heat-drought-paper).

GAPIT was run with default options, with the addition of the “Multiple_analyses = TRUE” option to output combined Manhattan plots and quantile-quantile (Q-Q) plots for each run (Wang and Zhang, 2021). A kinship matrix was created using the GAPIT VanRaden method (VanRaden, 2008; Wang and Zhang, 2021). All other outputs are standard from GAPIT. The GWAS results were analyzed using ZBrowse (Ziegler et al., 2015), an interactive GWAS viewer that runs on R (R Core Team, 2020). The GWAS results from GAPIT were uploaded to ZBrowse and aligned to the most recent *B. distachyon* reference genome (version 3.2 from 2020) from Phytozome (Goodstein et al., 2012). The two genes closest to each significant SNP (one upstream and one downstream) from each GWAS were examined.

For more detail about the data analysis, scripts for all analyses done in R are provided in the archived GitHub repository (https://github.com/danforthcenter/brachypodium-heat-drought-paper).

## Author Contributions

MAG, EL, and TCM conceived the research; MAG designed the experiments; MAG and NF developed tools for the experiments; EA, KH, KG, MAG, TF, performed the experiments; EL, MAG, JB, NF, SP, JS analyzed images and data; EL and MAG wrote the article; EL, JB, JS, KG, MAG, NF, edited the article.

## Acknowledgements

We would like to thank Kevin Reilly and his team for growth chamber and greenhouse plant care and support. We would also like to thank Melinda Wilson for expert care of the Bellwether Phenotyping Platform during the course of these experiments, to the Bellwether Foundation for the generous donation that allowed for the construction of the Bellwether Phenotyping Platform, and DDPSC Facilities for careful maintenance of the Bellwether Platform.

## Figure Legends

***Supplemental Figure 1 (collection location climate data table):***

Table of all 149 accessions included in this study with collection location and climate of origin data. Collection location information (Country, Longitude, Latitude, Elevation) from original collectors, and climate data from WorldClim v2.1 (Fick and Hijmans, 2017).

***Supplemental Figure 2 (table of all ranked phenotype results):***

Table of accessions ranked in descending order of performance in each treatment for the three main phenotypes assessed (height, area, percent damage).

***Supplemental Figure 3 (area heatmaps):***

Heatmaps of the average plant shoot area in the four experimental conditions (from left to right: control, drought, heat, heat & drought combined). This was calculated by taking the mean of the area of all plants of the same accession in the same treatment in each imaging cycle. The x-axis of each heatmap shows “Imaging Day,” which represents each imaging cycle, since half of the plants were imaged on even days and half on odd (so Imaging Day 1 is all images taken on day 1 *and* day 2 of the experiment). Imaging Day 2 was removed from the control and drought conditions, and Imaging Day 3 was removed from the heat treatment. On the y-axis are the accessions, sorted alphabetically in descending order.

***Supplemental Figure 4 (tolerant/susceptible accessions table):***

Table of accessions identified as being either tolerant or susceptible to the drought, heat or combination drought and heat stress. Accessions were identified as being tolerant when they were in the top 25% of all accessions for all of the three main phenotypes assessed (height, area, percent damage) for a given treatment, and susceptible when they were in the bottom 25% for all three phenotypes for a given treatment. Accessions were assessed by taking the difference between the measurement in the stress treatment and control conditions and comparing this difference between accessions.

***Supplemental Figure 5 (scree plot):***

Scree plot of the principal components (PC) from the principal component analysis (PCA) conducted with the climate of origin data. This plot shows the PCs on the x-axis and the percentage of variance explained by the PCs on the y-axis. Higher y-values indicate more variance explained by that PC.

***Supplemental Figure 6 (table of loadings):***

Table of loadings of the PCs in the PCA of the climate of origin data. This table shows how the bioclimatic variables contribute to each of the PCs. All 20 variables contribute to each PC, but values greater than |0. 2236| are considered significant and are bolded. Variables are colored by variable type (red for temperature-related, blue for precipitation-related, purple for both temperature- and precipitation-related, and white for elevation), and are roughly sorted in descending order of contribution to PCs.

***Supplemental Figure 7 (A: area Manhattan plots, B: percent damage manhattan plots):***

Manhattan plot of GWAS discovery results for plant shoot area (**A**) and percent damage (**B**). Each point represents a SNP. The height of the SNP on the y-axis represents the strength of association with plant shoot area (**A**) or percent damage (**B**), expressed as −log10(p-value). Points are colored by chromosome, and appear darker and more solid with lower p-values (stronger association with plant trait). The green horizontal line represents the significance cutoff and is calculated by dividing 0.05 by the number of SNPs included in the GWAS: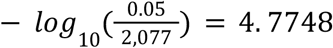

***Supplemental Figure 8 (table of significant SNPs):***

Table of SNPs found by GWAS to be significantly associated with height, area, or percent damage with information about genes close to these SNPs. Includes information about SNPs, genes closest to SNP on either side, and potential function of these genes.

***Supplemental Figure 9 (A: LD decay plot, B: LD decay values):***

**A**) Plot representing linkage disequilibrium (LD) decay for this *B. distachyon* population. The x-axis represents distance apart in base pairs (bp), and the y-axis is LD, based on the r^2^ value.

