## Supplementary figures and images for "Natural Variation in *Brachypodium distachyon* Responses to Combined Abiotic Stresses"

### Supplemental Figure 3

#### Supplemental Figure 3

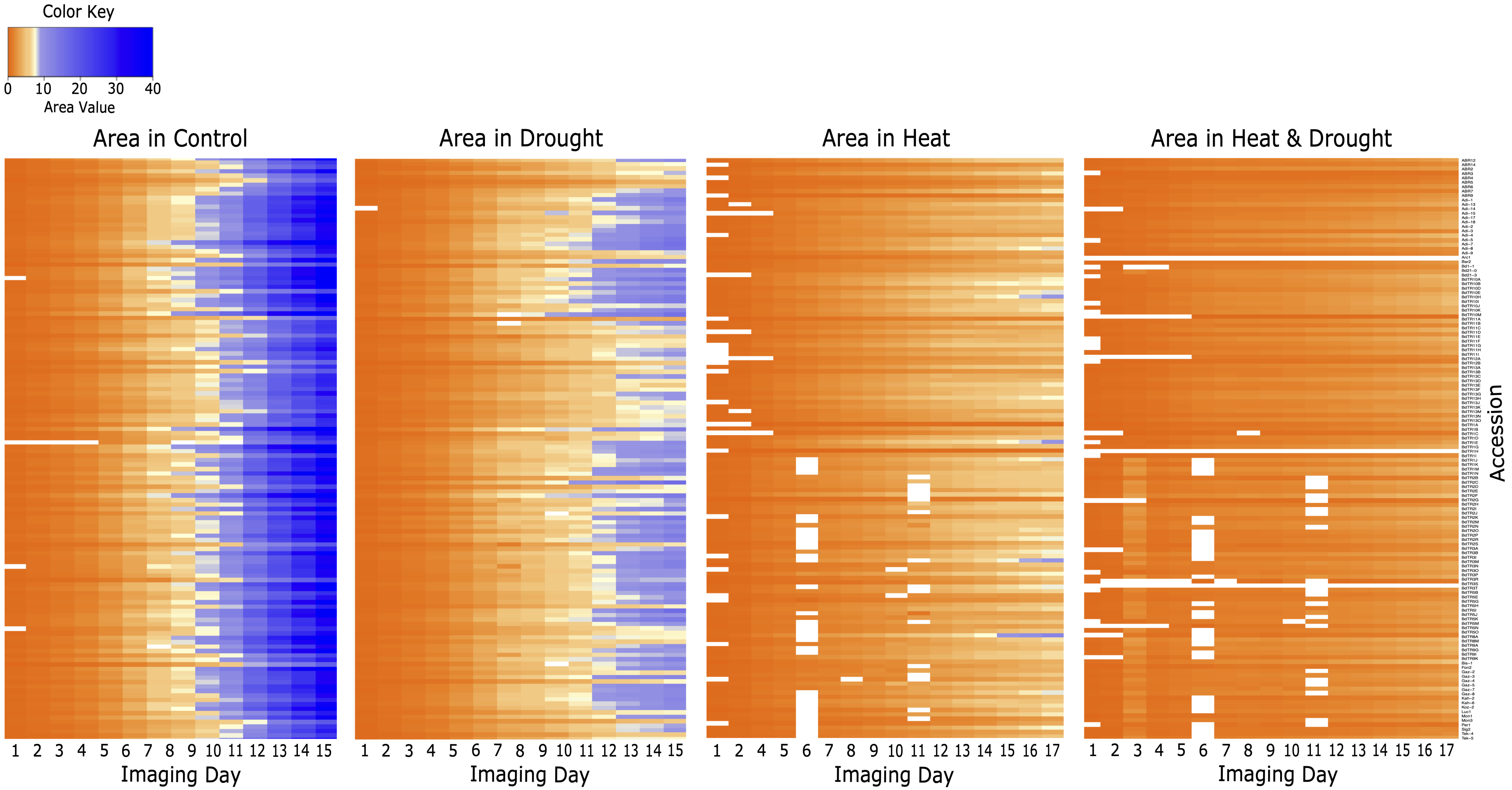

### Supplemental Figure 5

#### Supplemental Figure 5

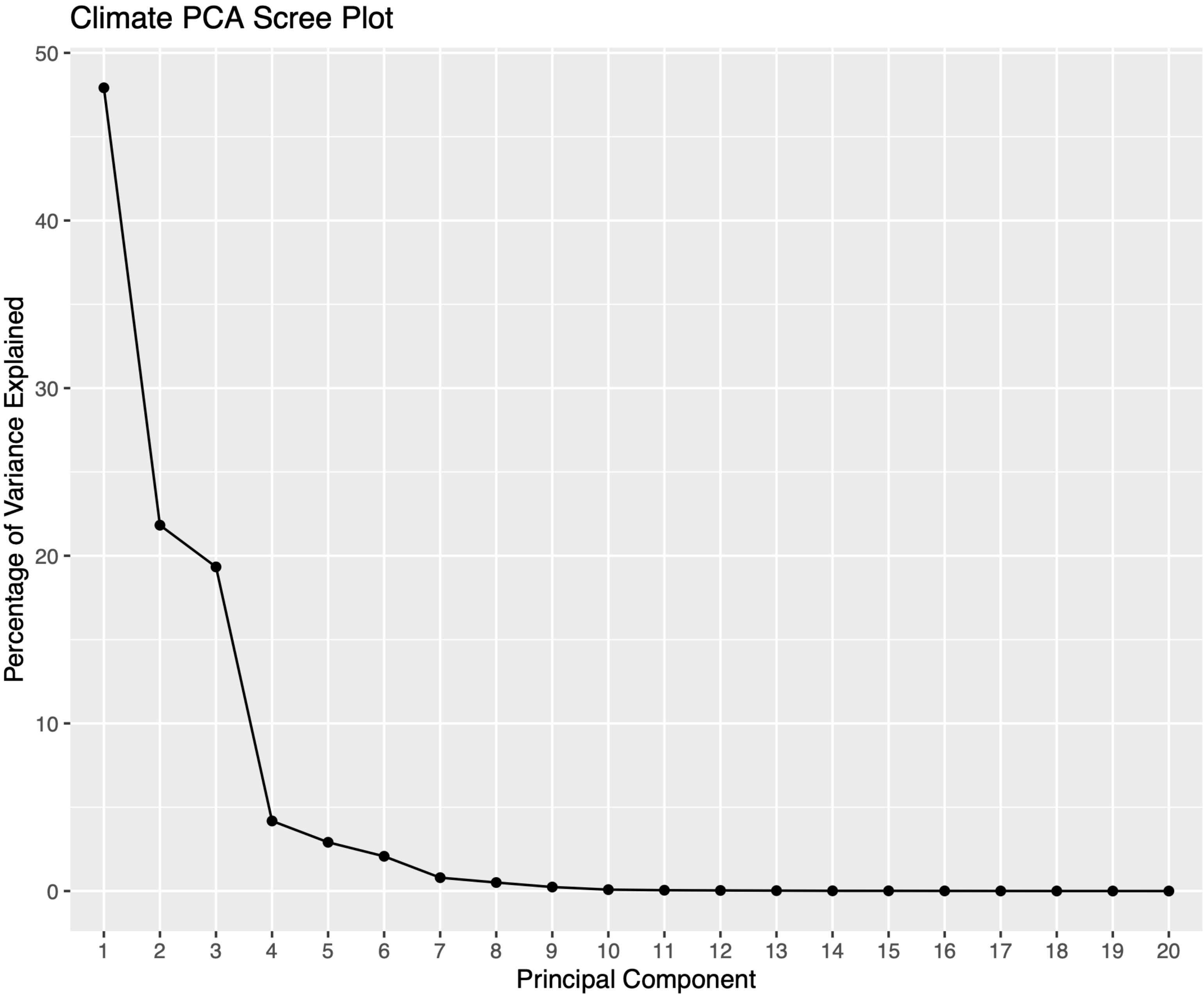

### Supplemental Figure 7

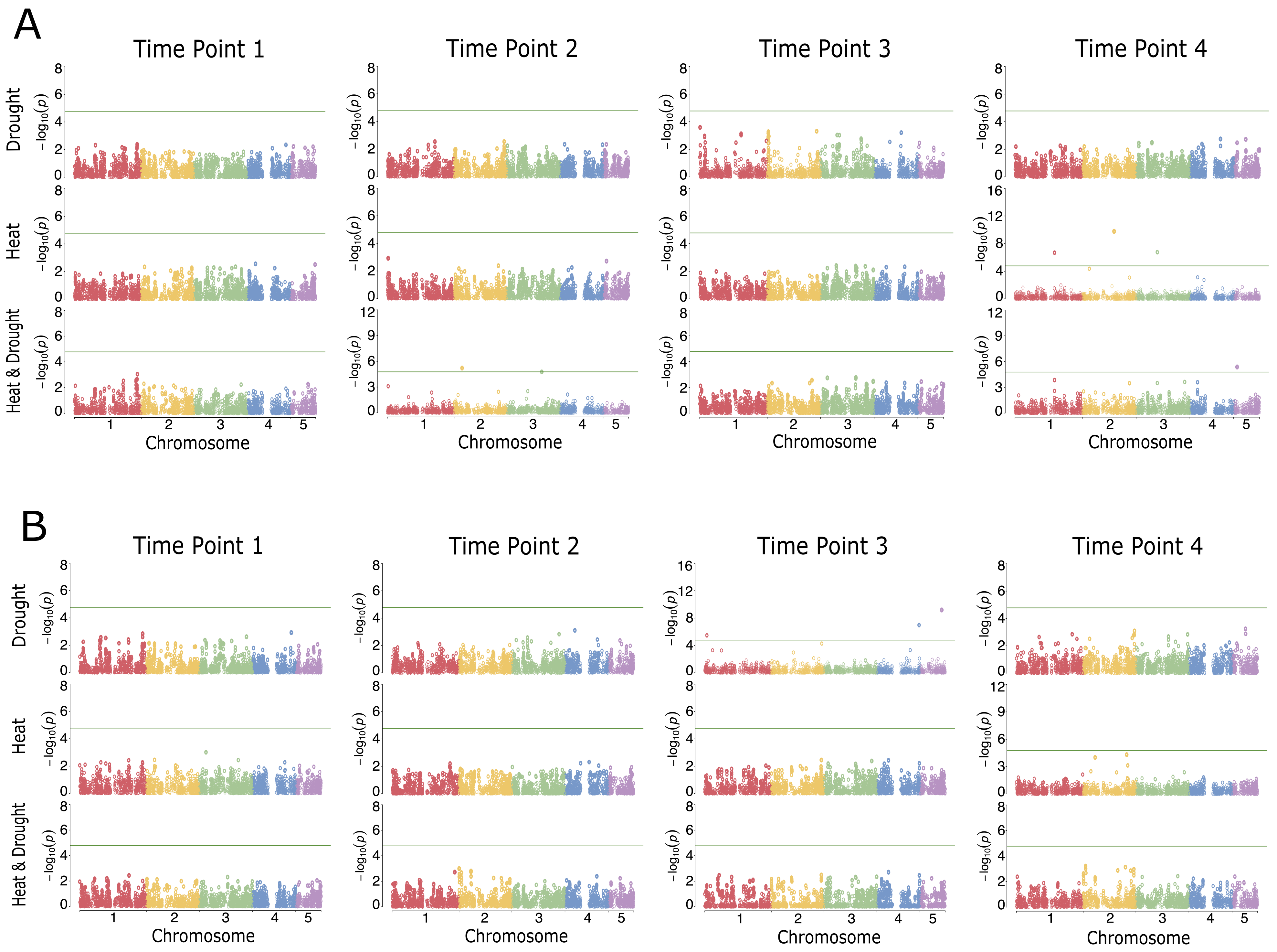

### Supplemental Figure 9

#### Supplemental Figure 9

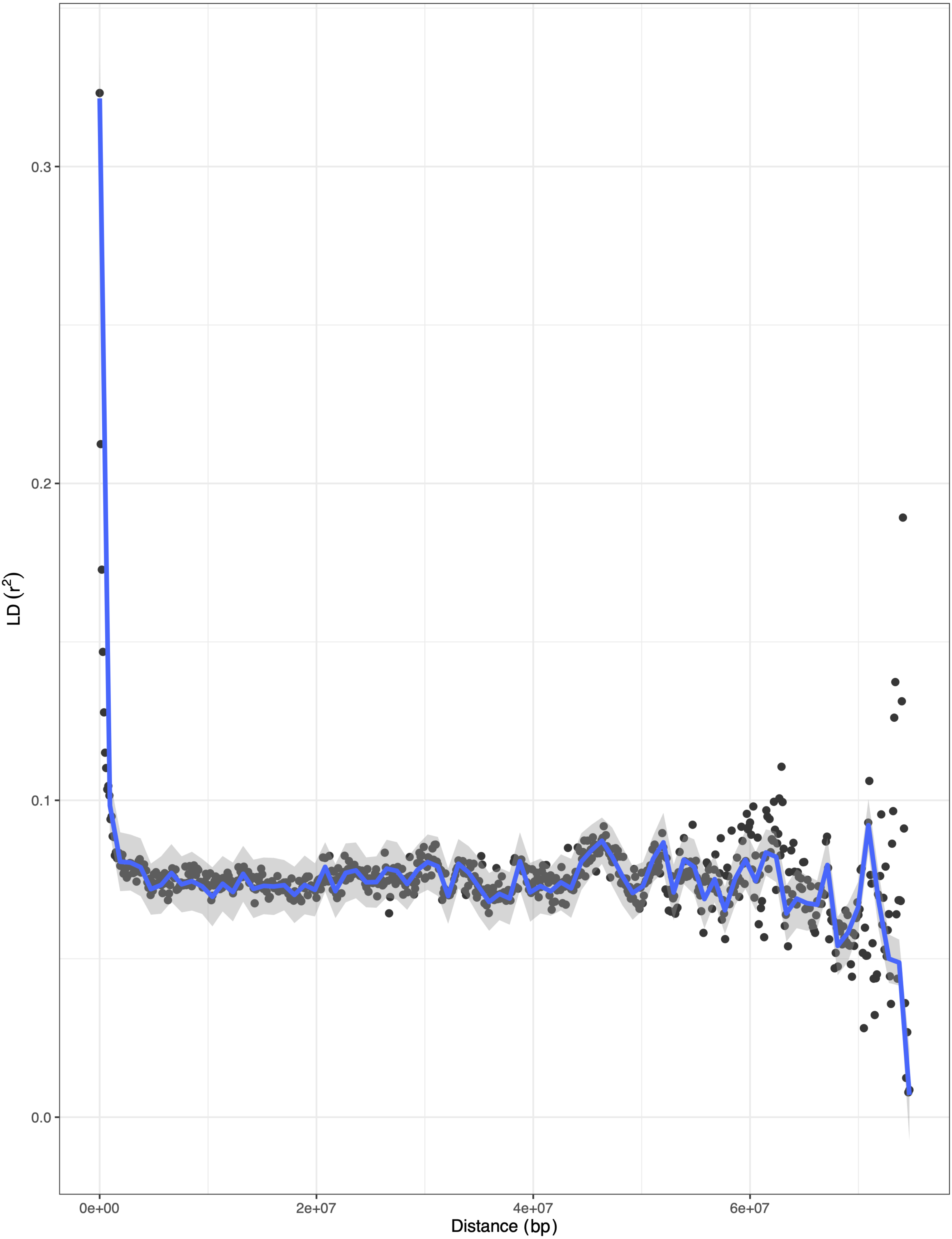

### Supplemental Figure 9A

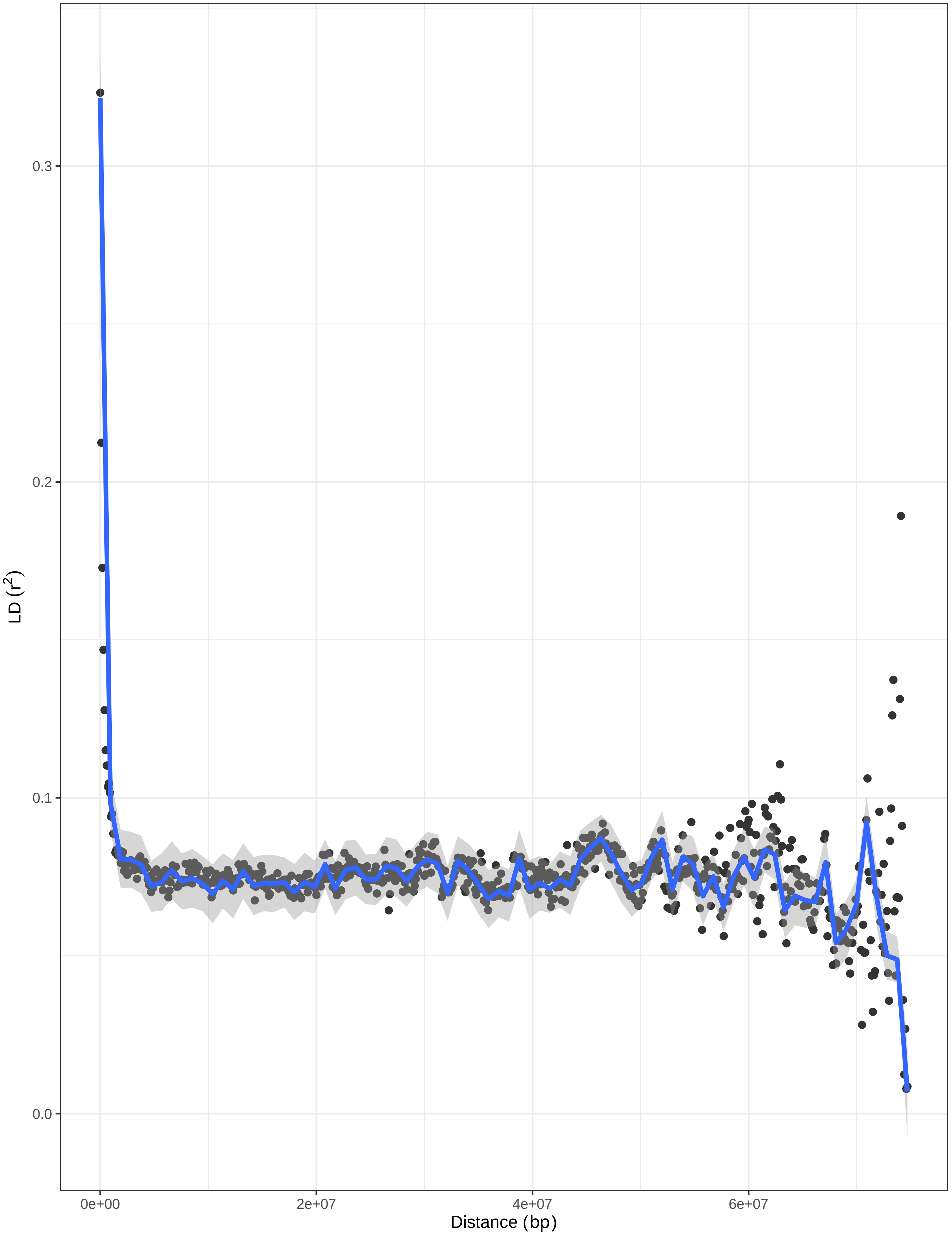
